# RareFold: Structure Prediction and Design of Proteins with Noncanonical Amino Acids

**DOI:** 10.1101/2025.05.19.654846

**Authors:** Qiuzhen Li, Diandra Daumiller, Fanglei Zuo, Harold Marcotte, Qiang Pan-Hammarström, Patrick Bryant

## Abstract

Protein structure prediction and design have traditionally been confined to the 20 canonical amino acids. Expanding this chemical space to include non-canonical amino acids (ncAAs) is essential for engineering proteins with novel chemical and functional properties. However, existing methods are not designed to generalise across chemically diverse residue types. Here, we present RareFold, a deep learning architecture for structure prediction and design of proteins containing the 20 canonical amino acids and 29 ncAAs. By representing each residue as an independent token, RareFold learns context-dependent atomic interaction patterns across chemically diverse sequence spaces, enabling modelling of non-standard chemistries within a unified framework. We apply this capability in EvoBindRare, a generative framework for de novo design of linear and cyclic peptide binders with an efficient implementation that substantially reduces computational requirements compared to existing architectures. We demonstrate its performance by designing binders against Ribonuclease A, yielding novel linear and cyclic peptides incorporating ncAAs within predicted interfaces with low-micromolar affinities (K_D_ ∼2-9 μM), comparable to the native ligand (K_D_ ∼2 μM). Hydrogen-deuterium exchange mass spectrometry confirms that the designed peptides engage the target at regions consistent with predicted binding interfaces. In addition, immunogenicity profiling in human-derived organoid models shows no detectable immune activation. By extending deep learning-based protein design to non-canonical chemical spaces, RareFold enables programmable access to expanded amino acid alphabets and broadens the scope of *de novo* protein engineering.

## Introduction

Structure prediction with AlphaFold2 (AF2) has revolutionised structural biology by achieving near-experimental accuracy [1], enabling new possibilities in rational protein engineering. Beyond predicting natural protein structures, structure prediction now plays a central role in *inverse design*, where new sequences are generated to adopt desired structures and functions [2]. Most successful design pipelines [3,4] rely on AF2 or similar models to evaluate whether candidate sequences fold as intended [4–6], making forward prediction a key step in modern protein design.

However, these approaches are limited by the chemistry of the 20 canonical amino acids shared across life, restricting the range of possible interactions, stability, and functionality. While cyclic peptide design can improve stability [7–9], expanding to noncanonical amino acids (ncAAs) offers far greater potential. Indeed, the clinical and commercial validation of platforms such as the RaPid system [10] and the development of Merck’s oral PCSK9 inhibitor (MK-0616) [11] underscores the transformative power of expanding the amino acid vocabulary beyond nature’s standard set. Over 300 amino acid types are found in the Protein Data Bank (PDB), and more than 500 occur in nature [12], with 140 ncAAs known to be naturally incorporated into proteins [13,14]. Despite this, ncAAs remain largely unexplored in design. They offer unique therapeutic benefits [15], including enhanced protease resistance [16] and reduced immunogenicity [17] by escaping standard immune surveillance mechanisms. Furthermore, ncAAs significantly expand the catalytic repertoire of proteins, enabling novel functionalities in catalysis [18] and transition metal coordination for artificial metalloenzyme design [19].

There are two main challenges in predicting structures with ncAAs. First, standard multiple sequence alignments (MSAs) cannot distinguish these residues, as most appear as “X” in sequence databases, masking their identity. Nonetheless, the presence of “X” still conveys useful information about variability and alignment context. Second, while recent frameworks like AlphaFold3 (AF3) [20] can model modified residues atom by atom, it represents ncAAs by mapping them to the closest canonical amino acid. Consequently, specifying each modified atom is impractical for design workflows due to the vast search space and changing input/output dimensions. In addition, it is not known how accurate AF3 is in predicting different types of modified amino acids, especially on unseen proteins, as it has been trained on almost all proteins, making evaluation difficult.

To overcome these limitations, we introduce RareFold, a structure prediction network trained on the standard 20 amino acids plus 29 additional ncAAs. By representing each residue, canonical or modified, as a unique token, RareFold learns coevolutionary and structural relationships in a compact, tractable manner. This representation enables both accurate structure prediction and joint sequence-structure optimisation. We further demonstrate its power by inverting the network into a design framework, EvoBindRare, which successfully designs linear and cyclic peptide binders incorporating ncAAs.

## Results

### RareFold

The genetic code comprises 64 codons, but due to redundancy, it typically encodes only 20 standard amino acids in proteins. In contrast, nature employs a much wider chemical repertoire: over 500 amino acid types have been identified [12], and 140 of these noncanonical amino acids (ncAAs) are known to be incorporated into proteins [13], offering an expanded landscape for molecular interactions and functionality. The PDB contains 331 distinct amino acid types in single-chain proteins, underscoring the untapped potential of ncAAs in protein engineering.

To enable robust structure prediction across this chemically diverse space, we extended the EvoFormer architecture [1] to include 29 of the most common ncAAs found in the PDB. In contrast to previous efforts [20,21], which operate at the atomic level, our approach treats each amino acid, canonical or noncanonical, as a unique token. This design choice, in principle, allows the model to learn sequence-structure relationships that influence how ncAAs are incorporated into protein structures (Figure 1). RareFold is a structure prediction framework that explicitly represents and models rare ncAAs.

**Figure 1.**
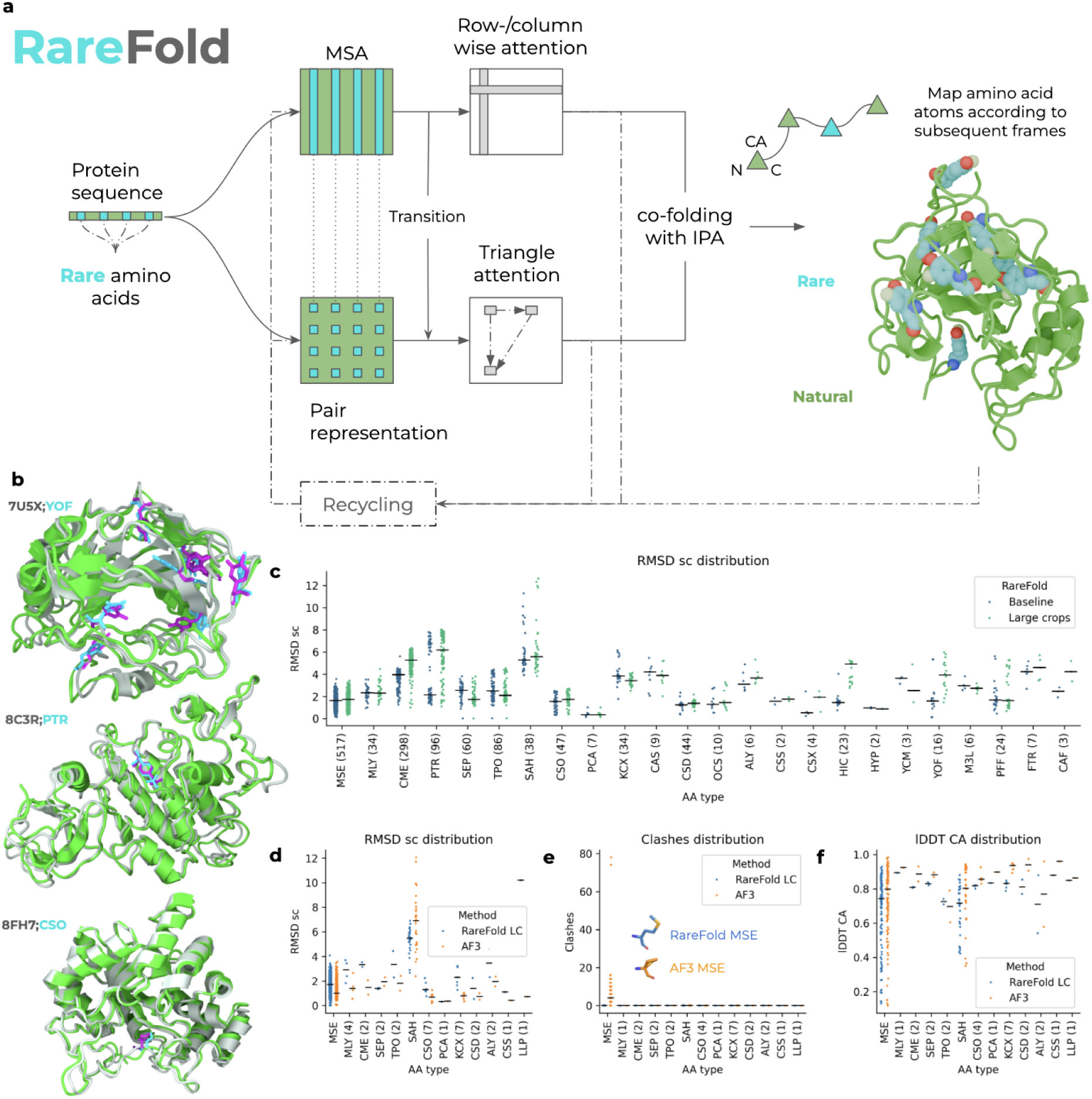
RareFold network architecture and evaluation. **a)** RareFold is a structure prediction network based on the EvoFormer architecture from AlphaFold2, trained to model both the 20 canonical amino acids and 29 additional noncanonical amino acids (ncAAs). Each amino acid, whether canonical or modified, is treated as a distinct token, enabling the model to learn residue-specific structural and coevolutionary patterns. This tokenised representation is essential for supporting iterative sequence optimisation and forward prediction in design tasks. For each input sequence, a multiple sequence alignment (MSA) is generated without templates; positions corresponding to ncAAs are marked as “X” due to missing homology information, shown in cyan. The MSA is processed using row and column attention, while pairwise residue interactions are captured using triangle attention mechanisms. Final residue representations are mapped to atomic coordinate frames, with each amino acid assigned a specific set of local frames, resulting in 49 distinct frame types in total. This architecture enables accurate structural modelling across chemically diverse protein sequences. **b)** Examples of accurate RareFold predictions across diverse folds and ncAAs using the baseline model. Superpositions of predicted (green cartoon) and native (grey cartoon) single-chain protein structures containing the ncAAs YOF (PDBID 7U5X), PTR (PDBID 8C3R), and CSO (PDBID 8FH7). The ncAA side chains are shown as sticks: RareFold predictions in cyan, and native PDB structures in magenta. These examples illustrate RareFold’s ability to accurately model structurally diverse proteins incorporating chemically distinct ncAAs. **c)** Distribution of aligned side chain RMSD (Å) on the validation set divided by ncAA type and number of recycles used at inference (3, 5, 10 and 20). The number of ncAAs evaluated is provided in parentheses next to each AA type. **d)** Distribution of aligned test set side chain RMSD (Å) per ncAA type for RareFold (LC=large crop mode, n=174) and AlphaFold3 (AF3, n=171). The number of ncAAs evaluated is provided in parentheses next to each AA type. There are two discrepancies since AF3 was OOM for some structures: MSE 118 vs 120 and SAH 31 vs 32, for AF3 and RareFold, respectively. **e)** Clash distribution within each ncAA summed per target on the test set. Clashes are defined as atoms within each ncAA closer than 1 Å apart. Only AF3 exhibits clashes for MSE, due to the side-chain being folded in on itself (see the visualisation). This is analysed in more detail in Supplementary Figure 2. The number of ncAAs evaluated is provided in parentheses next to each AA type. There are two discrepancies since AF3 was OOM for some structures: MSE 118 vs 120 and SAH 31 vs 32, for AF3 and RareFold, respectively. The number of structures is n=174 for RareFold and n=171 for AF3 (3 could not be predicted due to memory limitations). **f)** Test set Cα lDDT per protein grouped by the ncAA type the protein contains for RareFold (LC=large crop mode, n=174) and AlphaFold3 (AF3, n=171). The number of ncAAs evaluated is provided in parentheses next to each AA type. There are two discrepancies since AF3 was OOM for some structures: MSE 118 vs 120 and SAH 31 vs 32, for AF3 and RareFold, respectively

We evaluate RareFold on a held-out set of structures with less than 20% sequence identity to the training data. RareFold accurately predicts both the overall protein fold and the placement of diverse ncAAs (Figure 1b). In Figure 1c, we report the side chain (sc) RMSD for 24 different ncAA types across 731 validation structures, comparing the baseline RareFold model (trained on 256-residue crops) against a fine-tuned version (trained on 512-residue crops, Methods). The RMSD values are consistently low and comparable to those of standard amino acids (see *Comparison with proteinogenic amino acids*), indicating that RareFold generalises well across chemically diverse residues regardless of crop size. We observed a minor trade-off during fine-tuning, where larger crops improved global metrics but slightly increased side-chain RMSD for specific residues (e.g., CME, PTR). Notably, simply increasing the number of recycling iterations did not yield substantial improvements (Supplementary Figure 1), suggesting that the model achieves accurate predictions efficiently without requiring extensive inference-time compute.

In Figures 1d-f, we compare RareFold and AlphaFold3 (AF3) on a test set of 174 structures (171 for AF3 due to memory constraints), covering the 13 ncAA types remaining after strictly excluding AF3’s training data. Performance is largely comparable, with side chain RMSDs differing by only 1-2 Å for most amino acids. However, distinct failure modes emerge in the atomistic AF3 model. While AF3 ostensibly outperforms RareFold on MSE (Selenomethionine), this lower RMSD is an artefact of the diffusion process: AF3 frequently predicts physically impossible, collapsed geometries where all atoms overlap, minimising RMSD by placing atoms near the geometric mean but resulting in severe steric clashes. In contrast, RareFold generates physically valid, extended side-chain conformations for MSE (Figure 1e, Supplementary Figure 2). RareFold outperforms AF3 on residues such as SEP, SAH, and PCA. The case of S-adenosyl-L-homocysteine (SAH) is particularly notable; this residue is often not covalently linked in PDB structures, making its prediction akin to ligand docking [15]. While AF3 tends to treat SAH as covalently bound, potentially introducing errors, RareFold remains agnostic to covalent linkage, allowing for more accurate placement.

Overall, RareFold achieves high structural accuracy, reaching a median Cα lDDT of 0.96 on the validation set (Figure 2). When grouped by ncAA type on the test set (Figure 1f), the median Cα lDDT improves from 0.56 (baseline) to 0.76 after fine-tuning on large crops, approaching the 0.84 achieved by AF3 (Supplementary Figure 3). It is important to contextualise this gap: AF3 was trained on crops of 768 residues and utilised self-distillation on over 200 million structures predicted with AF2 (effectively seeing all known protein folds), whereas RareFold was trained on only ∼75,000 structures (0.04% of the AF3 training set). Despite this massive data disparity, RareFold demonstrates comparable capacity for modelling both canonical and ncAAs. We note that the few available structures for evaluating AF3 make it difficult to draw reliable conclusions about its ability for ncAA modelling outside MSE, where clashes are exhibited.

**Figure 2.**
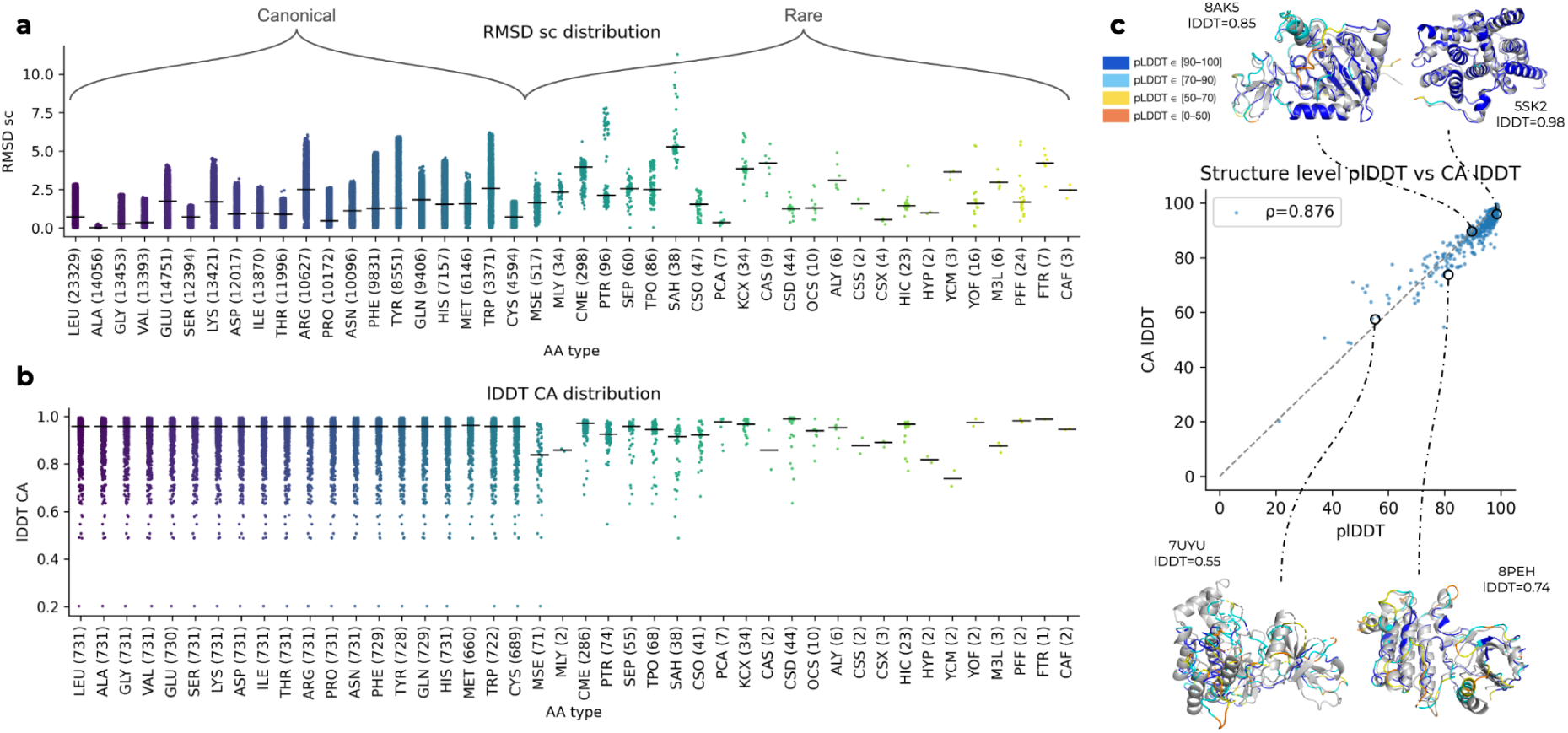
Accuracy of canonical and non-canonical amino acid modelling. **(a, b)** Performance comparison on the held-out validation set (n=731 structures; fine-tuned on 512-residue crops). **a)** Distribution of sidechain RMSD (Å) for canonical and non-canonical amino acids. The number of occurrences for each residue type is shown in parentheses. RareFold maintains low RMSD values for ncAAs comparable to canonical residues (e.g., CME vs. ARG), with higher variance observed only for rare types with limited training examples. **b)** Distribution of global Cα-lDDT scores. While ubiquitous canonical residues show consistently high median scores (>0.9), most ncAAs are also predicted with high accuracy, with performance drops limited to specific rare types (e.g., MLY, HYP). **c)** Correlation between the model’s predicted confidence (plDDT) and actual structural accuracy (Cα-lDDT) for full protein structures. The strong correlation (Spearman ρ = 0.876) indicates the model reliably self-assesses prediction quality. Insets display representative superpositions of predicted (coloured by plDDTl) and native (grey) structures across the accuracy spectrum, including 7UYU (lDDT=0.55), 8PEH (lDDT=0.74), 8AK5 (lDDT=0.85), and 5SK2 (lDDT=0.98), demonstrating that high-confidence predictions (plDDT >80) correspond to near-native backbone geometries.

Critically, RareFold’s token-based architecture offers a decisive advantage in computational efficiency. AF3 failed to process three test set structures even with 80 GB of vRAM, whereas RareFold predicted all structures using only 40 GB. This efficiency is transformative for protein design: by also batching predictions, RareFold enables a reduction in GPU memory footprint by 97% and accelerates the design cycle 6-fold (see *Methods*). This speed and low overhead enable the large-scale sequence-structure evaluations required for generative design, a capability currently impractical with current methods.

The ncAA side chains are shown as sticks: RareFold predictions in cyan, and native PDB structures in magenta. These examples illustrate RareFold’s ability to accurately model structurally diverse proteins incorporating chemically distinct ncAAs. **c)** Distribution of aligned side chain RMSD (Å) on the validation set divided by ncAA type and number of recycles used at inference (3, 5, 10 and 20). The number of ncAAs evaluated is provided in parentheses next to each AA type. **d)** Distribution of aligned test set side chain RMSD (Å) per ncAA type for RareFold (LC=large crop mode, n=174) and AlphaFold3 (AF3, n=171).

The number of ncAAs evaluated is provided in parentheses next to each AA type. There are two discrepancies since AF3 was OOM for some structures: MSE 118 vs 120 and SAH 31 vs 32, for AF3 and RareFold, respectively. **e)** Clash distribution within each ncAA summed per target on the test set. Clashes are defined as atoms within each ncAA closer than 1 Å apart. Only AF3 exhibits clashes for MSE, due to the side-chain being folded in on itself (see the visualisation). This is analysed in more detail in Supplementary Figure 2. The number of ncAAs evaluated is provided in parentheses next to each AA type. There are two discrepancies since AF3 was OOM for some structures: MSE 118 vs 120 and SAH 31 vs 32, for AF3 and RareFold, respectively. The number of structures is n=174 for RareFold and n=171 for AF3 (3 could not be predicted due to memory limitations). **f)** Test set Cα lDDT per protein grouped by the ncAA type the protein contains for RareFold (LC=large crop mode, n=174) and AlphaFold3 (AF3, n=171). The number of ncAAs evaluated is provided in parentheses next to each AA type. There are two discrepancies since AF3 was OOM for some structures: MSE 118 vs 120 and SAH 31 vs 32, for AF3 and RareFold, respectively.

### Amino acid accuracy across types and predicted confidence

Figure 2a presents the side-chain (sc) RMSD distribution for all amino acids in the validation set. We observe that predictive accuracy is comparable between canonical and non-canonical residues; for instance, Arginine (ARG) and the similarly sized CME both exhibit RMSD values spanning 1-5 Å. This variability captures the inherent structural uncertainty in protein prediction, as consistent sub-Ångström accuracy is often constrained by the experimental resolution of the training structures.

Many of the ncAAs do display higher median values than the proteinogenic ones, but this can be attributed to the big difference in their occurrence (e.g. CME occurs 298 times and ARG 10,627). The median global average Cα lDDT scores are also consistently higher for the proteinogenic AAs (Figure 2b). Again, this is related to their occurrence as the proteinogenic AAs are present in all structures, resulting in the overall median of 0.87, while the ncAAs are not. Still, many of the structures with ncAAs are accurately predicted and contain some examples with Cα lDDT >0.9 (all except MLY, SAH, CSS, CSX, HYP, YCM and M3L).

In design settings, it is important to both accurately predict structures and to identify the accurate predictions. The predicted lDDT score [22] (plDDT) provides a confidence measure for individual residues and the overall structure. As shown in Figure 2, we can reliably identify the overall structural accuracy, using plDDT (Figure 2b, Spearman ρ = 0.876, median Cα lDDT=95.8). Notably, the vast majority of predictions achieve lDDT scores above 80, indicating high structural quality. Figure 2c highlights examples of predictions with high and low lDDT. For many ncAA types, the overall structure and atomic side chain details are both correctly predicted and identified by RareFold. At lDDT scores above 0.8, the predicted structures are almost identical to the native ones.

### Peptide binder design with noncanonical amino acids

The best utility of protein structure prediction is arguably protein design, as made accurate by AlphaFold2 [1]. It is possible to design single-chain proteins by inverting structure prediction networks; however, this does not necessarily result in functional outcomes. A more interesting, albeit difficult, problem is protein binder design - especially that of small binders, as these have the potential to traverse cell membranes and be administered orally. To test the ability of RareFold, *which has not seen any protein complexes during training*, for design, we invert the network and adapt it to design linear and cyclic peptide binders from sequence information, as in our previous work with the standard 20 amino acids (EvoBind [9]). We incorporate the 29 ncAA of RareFold into a new framework called EvoBindRare (EBR, Figure 3). Twelve different ncAAs are available from our supplier (MSE, MLY, PTR, SEP, TPO, MLZ, ALY, HIC, HYP, M3L, PFF and MHO), and we design both linear and cyclic binders using these and the canonical 20 AAs (32 AAs in total).

**Figure 3.**
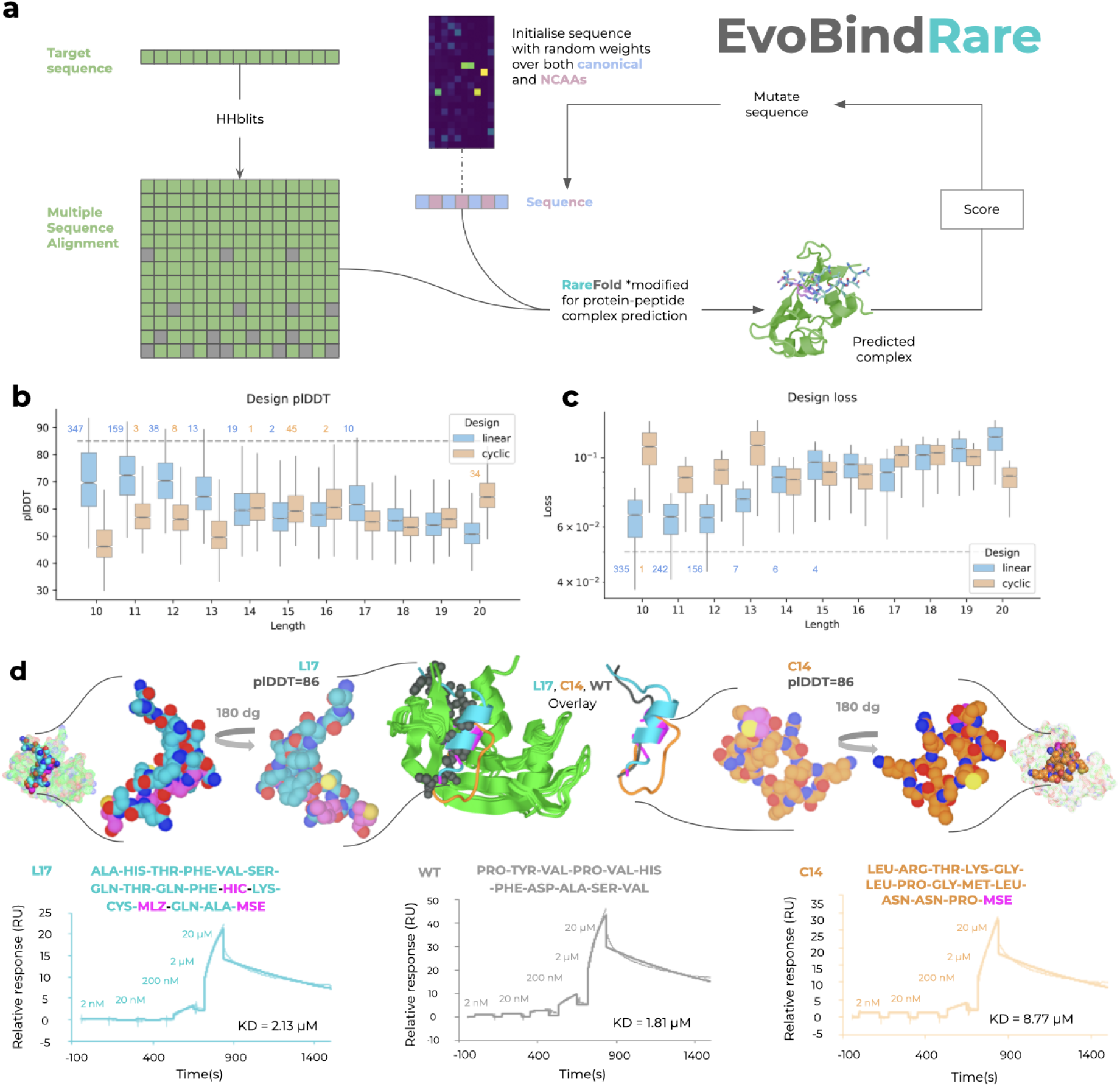
EvoBindRare design and experimental affinity measurements. **a)** Design of peptide binders using EvoBindRare (EBR). Starting from the sequence of a protein target, an MSA is generated with HHblits and a random peptide sequence is initialised. Both are input to a modified RareFold model for protein-peptide complex prediction. The predicted complex is scored (Methods), a mutation is introduced, and the process is repeated. EBR performs joint structure and sequence optimisation over 1000 mutation steps to design both linear or cyclic binders of 10-20 residues using up to 49 amino acid types. In this study, 32 amino acids (20 canonical + 12 ncAAs) were used due to synthesis constraints. **(b&c)** Optimisation results using EvoBindRare. The top 10% of designs, ranked by loss **(b)** and plDDT **(c)**, are plotted by peptide length (n = 4804 per length), with the hues indicating linear or cyclic topology. The dashed grey lines represent loss and plDDT cutoffs at 0.05 and 85, respectively. The number of designs satisfying these thresholds is annotated at each peptide length for both linear and cyclic scaffolds. To ensure visual clarity, outliers are excluded from the plot. These cutoffs reflect the primary design objective: to identify at least one high-quality binder per length. The distributions quantify the probability of success; the centre line marks the median, the box limits represent the interquartile range (IQR), and the whiskers extend to the non-outlier extremes. **d)** Predicted structures and single-cycle kinetics experiments obtained from surface plasmon resonance (SPR). The designed peptides are shown as spheres bound to the target individually, and in overlay with the WT bound to the target (cartoon). The sensorgrams show 5 injections from peptide concentrations 2 nM, 20 nM, 200 nM, 2 µM, and 20 µM. Both the linear (L17, cyan, Kd=2.13 μM) and cyclic (C14, orange, Kd=8.77 μM) designed peptides are predicted to bind to the same site on the target (green) with ncAAs (magenta) in the interfaces - although in different ways. The affinity measurements reveal that our design strategy successfully creates binders in the same micromolar affinity range as the WT peptide (Kd=1.81 μM, grey) towards the Ribonuclease target protein.

Figure 3a shows the design procedure with EBR towards a Ribonuclease (PDB ID 1SSC). Starting from the protein target sequence, we generate an MSA with HHblits [23], and the binding sequence is randomly initiated. The MSA and the binder sequence are then input to a version of RareFold modified for protein-peptide structure prediction. We use the model from step 20000, fine-tuned for 5000 additional training steps to deter inter-residue clashes (Methods). We run 1000 steps of mutation and design binders of different lengths (10-20 residues).

The top 10% of designs ranked by loss and their corresponding plDDT scores across peptide lengths, for both linear and cyclic cases, are shown in Figures 3b and c. While plDDT scores generally increase with length in the linear designs, this trend is absent in the cyclic ones, likely due to the structural constraints imposed by cyclisation and the resulting variability in binding modes. Horizontal lines at plDDT=85 and loss=0.05 indicate the thresholds for high-confidence structure and low design loss, respectively. These benchmarks reflect the true objective: to obtain at least one high-quality binder per length. The distributions themselves quantify the likelihood of achieving such a design. We selected designs with plDDT>85 and chose the lowest-loss sequence at each length for synthesis, resulting in 7 linear and 6 cyclic peptides. Only one of the cyclic peptides could be synthesised at high purity.

The binding affinities of the synthesised peptides were evaluated using surface plasmon resonance (SPR). Of the linear designs, one out of seven (14%) showed binding in the micromolar range, while the single tested cyclic peptide (1/1) also exhibited micromolar affinity. The linear (L17) and cyclic (C14) binders had affinities (Kd) of 2.13 μM and 8.77 μM, respectively, compared to 1.81 μM for the known wild-type (WT) binder (Figure 3d, see Supplementary Figure 4 for all sensorgrams). The successful linear binder contains three types of noncanonical amino acids (ncAAs), all predicted to interact with the target interface, while the cyclic binder includes one ncAA, also located at the predicted interface (Figure 3d). These results demonstrate that RareFold-enabled design can incorporate ncAAs into functional interfaces, yielding binders with experimentally confirmed affinity. Notably, the predicted binding modes and novel sequences indicate that EBR can generate novel interfaces with comparable affinity to the WT, without prior exposure to any protein complex. When aligning the sequences with Clustal Omega [24], one can see that there are only two matching residues between the WT and L17, with a gap introduced, while C14 has no matches (Supplementary Figure 5).

### Mapping of designed interfaces with HDX-MS

To assess whether the designed peptides engage the target pocket at the predicted interface, we employed Hydrogen-Deuterium Exchange Mass Spectrometry (HDX-MS). By monitoring the exchange rate of backbone amide hydrogens with solvent deuterium, we mapped the “protection footprint”, which reflects changes in solvent accessibility and backbone dynamics for our non-canonical designs (Linear L17 and Cyclic C14) relative to the wild-type (WT) ligand. It is important to note that the spatial resolution of HDX-MS is determined by the length of the proteolytic peptides obtained during digestion. Because deuterium uptake is averaged across the entire length of a detected peptide “read”, observed protection may reflect a strong interaction at a specific distal site that propagates across the fragment, rather than uniform shielding of the entire sequence. Consequently, we interpret these data as regional conformational constraints rather than single-residue contacts.

As illustrated in Figure 4, differential HDX analysis revealed that the WT ligand induces significant protection of distinct regions, corresponding to the expected peptide shielding of the binding pocket. Crucially, both the linear (L17) and cyclic (C14) designs elicited protection patterns in similar structural regions, confirming their engagement at the predicted interface. The observed variations in the protection profiles between the peptides suggest distinct binding dynamics and are consistent with the specific geometric differences in the predicted protein-peptide complexes. These regions of relative protection are quantitatively detailed in Supplementary Figure 6.

**Figure 4.**
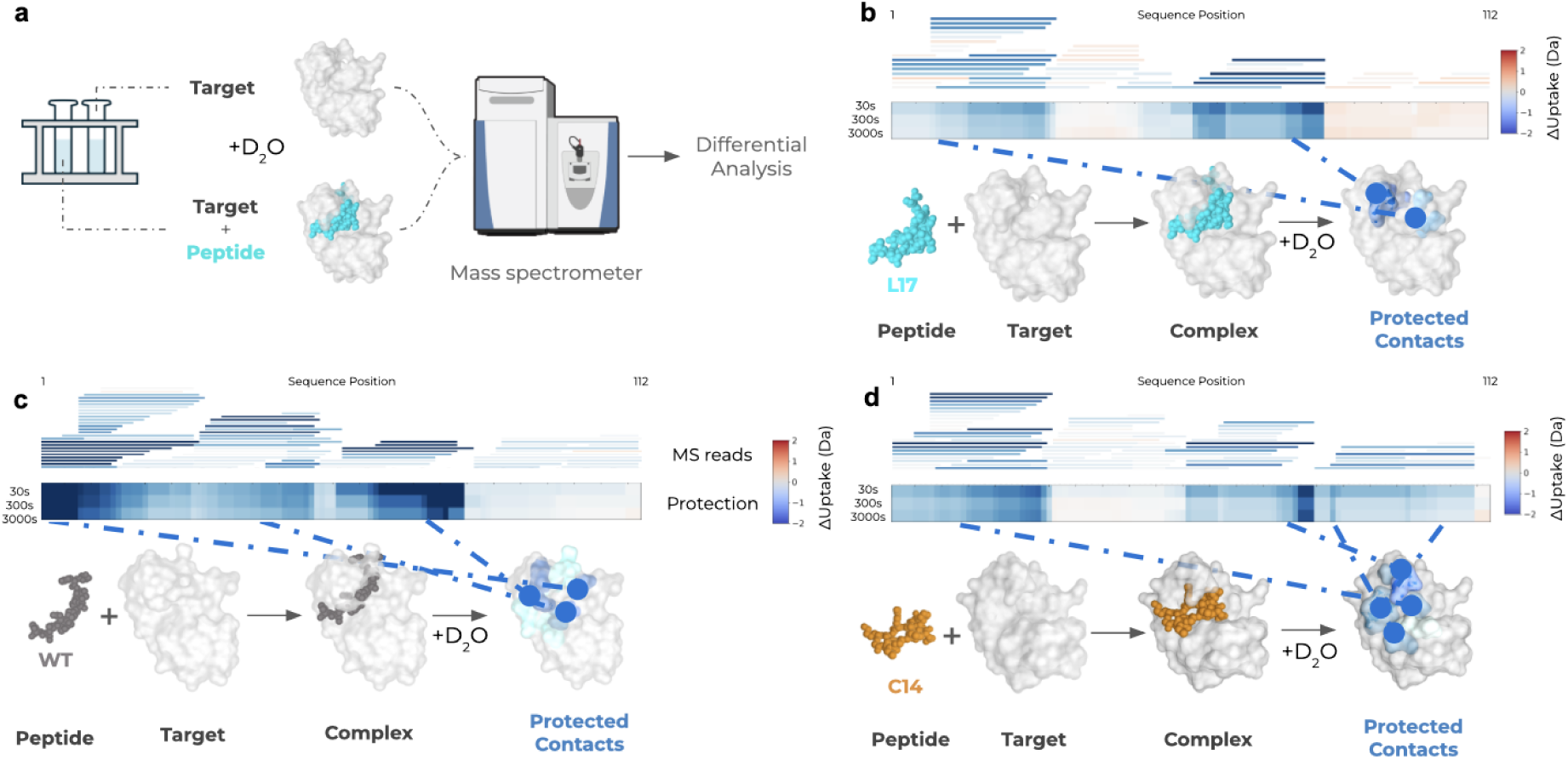
Differential HDX-MS mapping of peptide-target interfaces. **a)** To map the binding epitopes, the target protein (Ribonuclease A) was incubated with either the wild-type (WT) or de novo designed peptides (L17, C14) before labelling with deuterated solvent (D_2_O). Deuterium uptake was measured at 30s, 300s, and 3000s and compared against the uptake of the unbound (apo) target along the target length (1-112 residues). **(b-d)** Differential uptake analysis for **(b)** the WT peptide, **(c)** the linear design L17, and **(d)** the cyclic design C14. **Top panels (MS reads)**: Horizontal bars represent the proteolytic peptide fragments generated during digestion. Each bar is colored by the difference in deuterium uptake (Da) relative to the no-peptide control. **Middle panels (Heatmap)**: Aggregated protection data across the protein sequence over time. **Bottom panels (Structure)**: Surface representations of the predicted complexes. The target surface (predicted peptide contacts, residues with any atoms within 4 Å distance from the receptor, for L17 and C14, and structurally determined ones for the WT) is coloured according to the experimental HDX protection levels, visualising the binding footprint and conformational dynamics in solution. **Colour Scale**: The blue-white-red gradient indicates relative deuterium uptake (Da). Blue values indicate negative relative uptake (protection), where the bound peptide shields the target or stabilises the structure. Red values indicate increased uptake, reflecting conformational destabilisation or dynamic exposure due to binding/unbinding events. **Results**: While the linear (L17) and cyclic (C14) binders exhibit protection footprints that differ from each other and the WT, these distinct patterns align with the specific contact interfaces observed in their respective predicted structures. This confirms that the designs engage the target as predicted through the unique binding modes generated.

### Immunogenicity of peptides with ncAAs

Amino acid modifications, such as cyclisation, stapling, PEGylation, and lipidation, can significantly enhance peptide stability and pharmacological performance; however, they may also disrupt immune processing and increase immunogenicity [25,26]. This duality underscores the necessity of pairing proactive chemical optimisation with rigorous, stage-wise immunogenicity evaluation, including for process-related impurities, to ensure therapeutic efficacy without compromising patient safety [27].

To evaluate the immunogenicity of peptides containing ncAAs, C14 and L17 peptides were incubated at concentrations of 0.25, 2.5, and 25 µM with peripheral blood mononuclear cells (PBMCs) or immune tonsil organoids, each obtained from a different donor (Figure 5a). Immune responses were assessed by measuring inflammatory cytokine secretion and peptide-specific antibody production via ELISA, and phenotypic changes in immune cell subsets using flow cytometry (Methods). Neither C14 nor L17 induced cytokine production (IL-6, TNF-α, IL-1β, IFN-γ, and IL-12) in culture supernatants collected on days 2, 5, and 7 beyond the levels observed with the wild-type (WT) peptide, which served as the negative control during the seven-day incubation period (Figure 5b). Furthermore, no peptide-specific antibodies (IgG, IgM, or IgA) against WT, C14, or L17 were detected in supernatants from PBMC and tonsil organoid cultures on day 7, nor in pooled sera from three healthy donors (Figure 5c). In addition, no major differences were observed in the proportions of T cell and B cell subsets, or in monocyte populations, between cultures stimulated with C14, L17, or WT peptides (Figure 5d). By contrast, R848 (a potent TLR7/TLR8 agonist) and anti-CD3/CD28 (T cell activators), used as positive controls, induced robust cytokine production and notable alterations in both T cell populations and monocytes.

**Figure 5.**
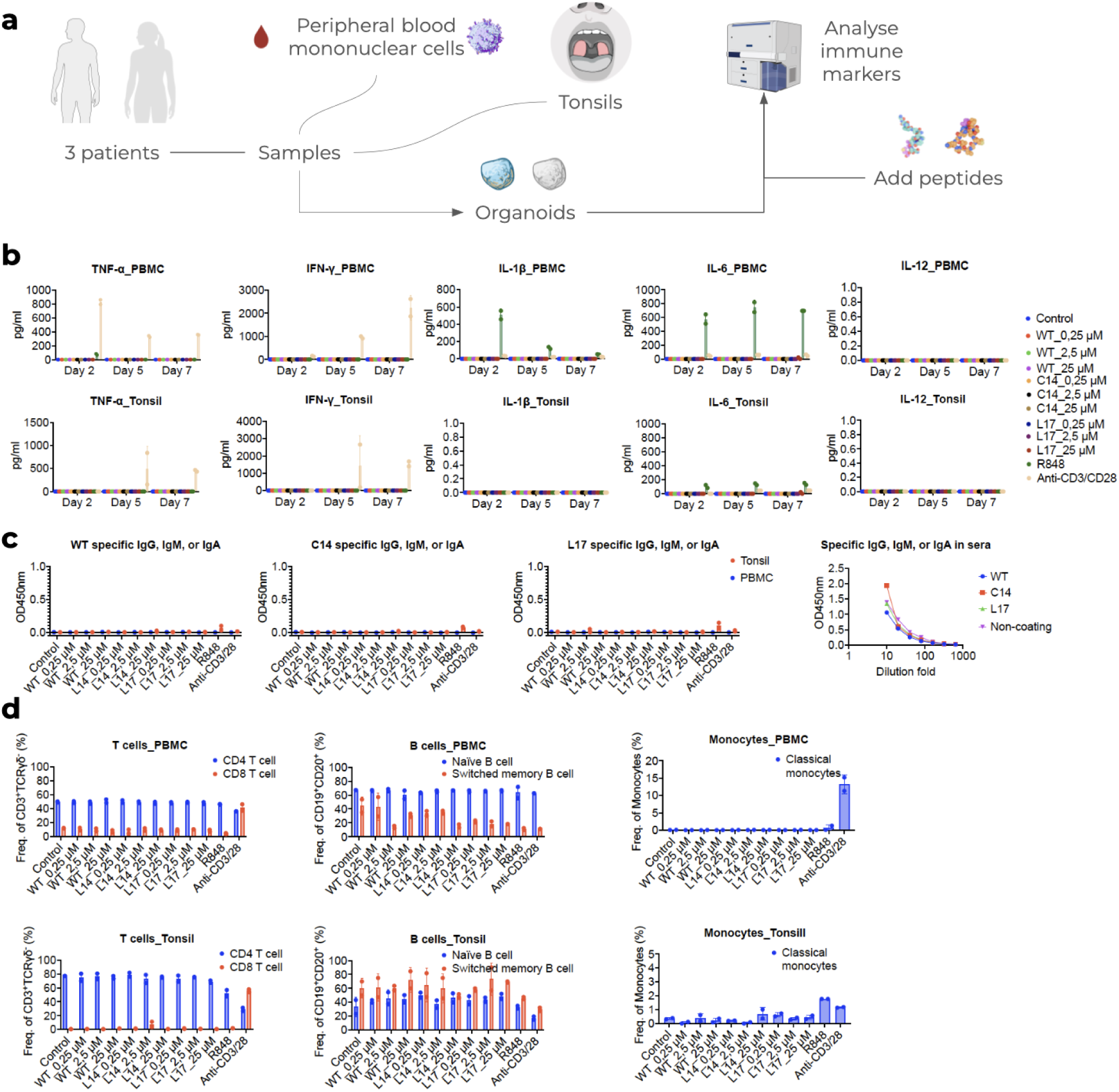
Immunogenicity of peptides containing ncAAs. **a)** PBMCs and tonsil organoids from independent donors were stimulated with WT, C14, or L17 peptides (0.25, 2.5, or 25 µM) for up to seven days. Each concentration was tested in duplicate wells. Positive controls were cells stimulated with the TLR7/8 agonist R848 and anti-CD3/CD28 antibodies. In all panels, the first column represents unstimulated cells. **b)** Cytokine concentrations (IL-6, TNF-α, IL-1β, IFN-γ, IL-12) in supernatants at days 2, 5, and 7, measured by ELISA (mean ± SD, pg/mL). C14 and L17 did not induce cytokine production above the WT baseline or control with no stimulation. R848 or anti-CD3/CD28 strongly induced IL-6, TNF-α, IL-1β, and IFN-γ, whereas IL-12 was undetectable under all conditions. **c)** Antibody responses assessed by ELISA. The first three panels show total peptide-specific antibody levels (IgG/IgM/IgA) in supernatants from PBMC and tonsil organoid cultures at day 7 (0.25–25 µM peptides), with unstimulated cultures as the control. The fourth panel shows pooled sera from three healthy donors tested across serial dilutions (1:20–1:640), with non-coated wells as the control. In all cases, OD450 values for peptide-coated wells (WT, C14, L17) did not exceed controls, indicating no peptide-specific antibodies. Neither R848 nor anti-CD3/CD28 induced antibody responses. **d)** Immune cell subsets analysed on day 7 by flow cytometry. Upper panels show PBMC cultures, and lower panels show tonsil organoid cultures. Left: CD4⁺ and CD8⁺ T cells as a proportion of CD3⁺γδ⁻ T cells. Middle: naïve (CD19⁺CD27⁻) and switched memory (CD19⁺CD27⁺IgD⁻) B cells as a percentage of CD19⁺CD20⁺ cells. Right: monocytes (CD14⁺, CD16⁺) as a frequency of total monocytes. No differences were observed between cell cultures stimulated with modified peptides and those stimulated with the WT peptide, whereas R848 and anti-CD3/CD28 induced alterations in T-cell distributions and monocyte frequencies.

## Discussion

The results show that RareFold enables accurate structure prediction of proteins incorporating a diverse set (29) of ncAAs, achieving performance comparable to AF3 on global structure and exceeding it on local side chain accuracy for some residues. Notably, RareFold produces more physically plausible structures for certain ncAAs, such as MSE, due to its frame mapping, avoiding intra-residue atomic clashes observed in AF3. While AF3 yields higher lDDT scores on average (0.84 vs 0.76), this is expected due to its training on a larger set of structures (including test-similar examples via self-distillation) and longer crop sizes. Although AlphaFold3 provides broad chemical coverage, it was not designed as a scalable inverse design framework. RareFold’s token-based representation enables direct integration into iterative design pipelines, allowing efficient inference even with limited computational resources. This efficiency is essential for practical protein design, where forward models are called repeatedly to explore large sequence-structure spaces. We show this by providing an efficient batched design implementation, allowing for using a single GPU in a design campaign, leading to a 97% reduction in memory consumption and 6 times faster design (Methods). This optimisation addresses the growing concern regarding the environmental impact of large-scale AI and democratizes access for smaller research groups with limited resources.Aa

Few existing forward models support design with ncAAs, and none, to our knowledge, have been systematically validated with both large-scale benchmarks and experimental tests. HighFold2, a recent transfer-learning extension of AlphaFold-multimer (AFM) [28], has attempted to include modified amino acids for cyclic peptide prediction (not design) [29]. Still, it is evaluated without separating structures seen during AFM pretraining, raising concerns about generalisability. Moreover, performance on liganded structures remains a major challenge, as evidenced by AF3’s results on novel interfaces and unseen targets with (10-20% success rate) [30]. A critical limitation lies in AF3’s implementation of modified residues: by necessitating that ncAAs be represented by their closest standard analogues in the Multiple Sequence Alignment (MSA), the model effectively discards that which define the ncAAs’ function. This approximation obscures the specific chemistry driving molecular recognition, where AF3 has previously been shown to fall short [31]. In design settings, this will likely lead to failures at complex interfaces where the modification itself, rather than the parent scaffold, dictates binding. These findings underscore the need for joint progress in computational methods combined with experimental evaluation. RareFold addresses this by offering a scalable and generalisable foundation, validated in the regime of experimental binder design, suggesting generalisation to novel tasks.

The EvoBindRare (EBR) platform is able to design both linear and cyclic peptide binders with ncAAs from scratch. The affinity to the ribonuclease target evaluated here is 2.13 μM and 8.77 μM, respectively, for the linear (L17) and cyclic (C14) binders, which is comparable to 1.81 μM for the known WT binder. When aligning the sequences, one can see that there are few matching residues between the designs and WT, demonstrating that the designs are not simple variants of the WT sequence (Supplementary Figure 5). Importantly, we do not observe a consistent increase in binding affinity relative to canonical designs. This is expected, as canonical amino acids are often sufficient for optimising affinity in well-characterised systems. However, the designed sequences incorporate multiple ncAAs while maintaining binding, indicating that these residues are compatible with functional interfaces and can be integrated without loss of activity. This supports the use of ncAAs to access chemical functionalities not present in the canonical set, including altered electrostatics, backbone constraints, and post-translational mimetics, which may be required for targets or design constraints where canonical residues are insufficient

While demonstrated here on the challenging, pH-sensitive Ribonuclease system, this success implies generalisability. Our original EvoBind [9] was similarly developed on this single target, yet successfully generalised to design inhibitors [32] and agonists [33] for unseen systems. Because the network has never seen protein complexes during training, its ability to generate binders with affinity comparable to WT stems from learning fundamental interface physics and coevolutionary constraints rather than memorising specific target motifs. We caution, however, that this generalizability remains to be empirically proven for non-canonical design, as the specific effects of ncAAs across diverse, novel target classes are currently unknown.

To validate the structural accuracy of these interactions beyond simple affinity measurements, we employed Hydrogen-Deuterium Exchange Mass Spectrometry (HDX-MS). While static methods like crystallography provide atomic snapshots, HDX-MS captures the dynamic engagement of the ligand in a physiological solution state. We observed that both the linear (L17) and cyclic (C14) designs induce protection footprints on the target protein that align with the predicted binding interfaces. Importantly, these protection patterns mirror those of the wild-type ligand in key functional regions. Although the spatial resolution of HDX-MS is limited by the length of the proteolytic peptides, meaning observed protection reflects regional conformational constraints rather than single-residue contacts, these data provide strong physical evidence that the de novo designs engage the target pocket as designed.

Immunogenicity profiling showed that C14 and L17 do not elicit higher responses than the WT. This aligns with prior reports that modifications such as D-amino acids or N-methylations can attenuate immune recognition by reducing MHC binding and T cell activation [34]. From a translational standpoint, these data suggest that peptides C14 and L17 may be suitable for long-term applications where minimising anti-drug antibody formation is essential. By contrast, structural interventions such as hydrocarbon stapling, β-amino acid incorporation, or cyclisation have been successfully applied to stabilise peptide conformations while intentionally enhancing immunogenicity for vaccines and cancer immunotherapy [25]. These findings underscore the importance of aligning chemical modification strategies with therapeutic goals and highlight the need for systematic *in vivo* validation and regulatory frameworks to balance pharmacological benefit with immunological safety [27]. Our work was based on *in vitro* PBMC and *ex vivo* tonsil organoid models with a single exposure over seven days. While these systems provide valuable insights into early innate and adaptive immune responses, they do not capture certain *in vivo* aspects, such as antigen biodistribution, repeated or long-term exposure, HLA diversity, or the influence of tissue-resident immune cells and the microenvironment. Complementary *in vivo* studies and longitudinal assessments would help to validate these findings.

The expansion from 20 to 49 amino acids results in a vast expansion in the sequence space (20^L^ vs 49^L^ for a protein of length L). However, extending this framework to the full spectrum of known ncAAs remains a significant challenge. The primary bottleneck is not architectural but data-driven: the scarcity of high-resolution structural data for rarer modifications precludes the creation of rigorous training and, crucially, held-out validation and test sets. Without ground-truth structures to benchmark against, the reliable inclusion of additional ncAAs would risk hallucination, undermining the physical validity that is central to RareFold’s utility. By uniting accurate ncAA structure prediction with efficient design strategies such as EBR, RareFold lays the groundwork for a new generation of protein design with expanded chemistry, provided that structural databases continue to grow to support the validation of these novel chemical spaces. The computational efficiency and zero-shot success of RareFold suggest its immediate utility for high-throughput therapeutic lead generation in industrial drug discovery pipelines.

## Methods

### Data

All monomeric protein structures from the PDB were selected on 2024-12-10, determined by X-RAY diffraction or Electron Microscopy (EM) with a resolution ≤5 Å (n=75’232 structures). We extracted the first protein chain in each PDB file and the corresponding sequences with less than 80% non-standard amino acids and more than 50 residues (n=74882/75232 proteins fulfilled these criteria, 99.5%). We clustered the sequences at 20% identity using MMseqs2 [35] with the command:

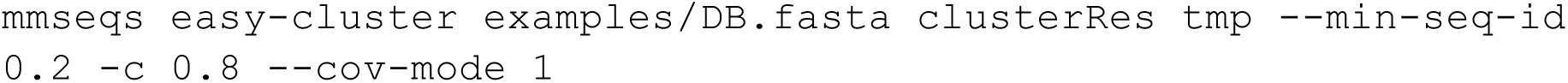

This analysis resulted in 9’031 clusters. We also examined the distribution of amino acid types across the 74’882 structures, which contained a total of 331 unique amino acids. However, the frequencies of ncAAs are much lower compared to the standard 20 amino acids commonly found in humans. Among the ncAAs, MSE is the most frequent, while the others occur with frequencies ranging from 10^1^-10^3^. To address the limited availability of data for ncAAs, we focus on learning from the top 50 amino acids by frequency, which includes 30 ncAAs. The rarest amino acid in this top 50 is MHO, appearing only 21 times (Figure 6).

**Figure 6.**
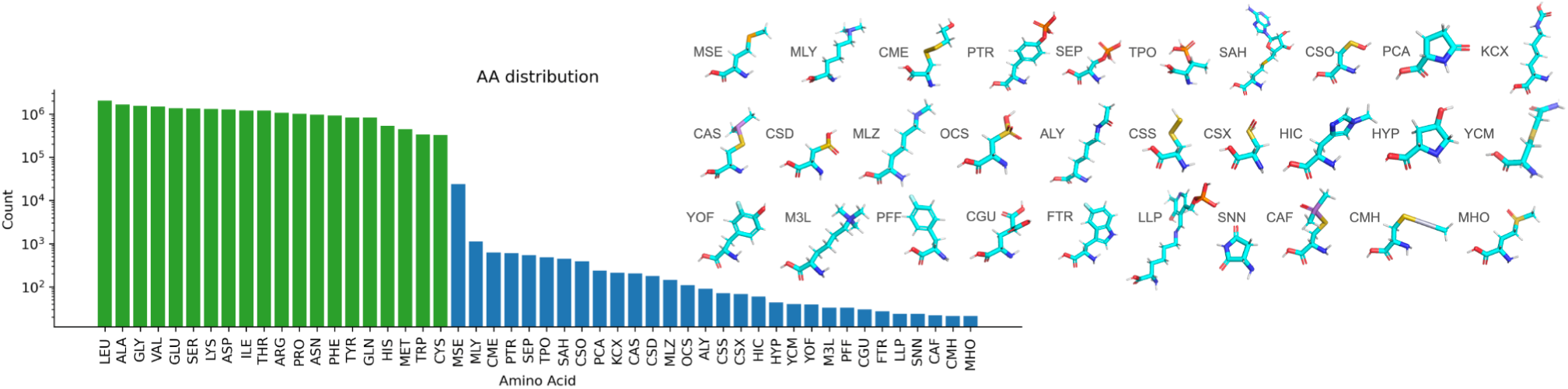
Distribution of amino acids in single-chain protein structures in the PDB. The distribution shows both proteinogenic (green) and noncanonical amino acids (ncAAs, blue) across single-chain protein structures. The 30 most common ncAAs (blue) were included in the training of RareFold. The minimum count in the top 50 most frequent amino acids is 21, represented by MHO. See Supplementary Figure 7 for the distribution of all amino acids present in the PDB.

### Noncanonical amino acid descriptions

To predict the coordinates of the additional 29 ncAAs (SNN was excluded since continuous peptide bonds can’t be formed with this amino acid) included in this study, we apply the following modifications to the frames available in AlphaFold2 for the standard 20 amino acids. New frames were implemented for atoms extending beyond the standard 20 amino acids structural frames available in AlphaFold2 (**Table 1**, see the code in the code availability section for a detailed implementation).

**Table 1:** Non-Canonical Amino Acids (ncAAs) in RareFold.

| Three-letter code | Description | Link |
| --- | --- | --- |
| MSE | S to Se in MET | https://www.rcsb.org/ligand/MSE |
| MLY | H to CH3 in LYS | https://www.rcsb.org/ligand/MLY |
| CME | H to S-C-C-OH in CYS | https://www.rcsb.org/ligand/CME |
| PTR | OH to PO4 in TYR | https://www.rcsb.org/ligand/PTR |
| SEP | OH to PO4 in SER | https://www.rcsb.org/ligand/SEP |
| TPO | OH to PO4 in THR | https://www.rcsb.org/ligand/TPO |
| SAH | add the large group to MET instead of the CH3 group as rigid | https://www.rcsb.org/ligand/SAH |
| CSO | H to OH in CYS | https://www.rcsb.org/ligand/CSO |
| PCA | add O to PRO | https://www.rcsb.org/ligand/PCA |
| KCX | add COO to LYS | https://www.rcsb.org/ligand/KCX |
| CAS | H to AS-C-C in CYS | https://www.rcsb.org/ligand/CAS |
| CSD | O2 to S rigid in CYS | https://www.rcsb.org/ligand/CSD |
| MLZ | H3 to CH3 in LYS | https://www.rcsb.org/ligand/MLZ |
| OCS | O3 to S in CYS | https://www.rcsb.org/ligand/OCS |
| ALY | CCO to N in LYS | https://www.rcsb.org/ligand/ALY |
| CSS | SH to S in CYS | https://www.rcsb.org/ligand/CSS |
| CSX | O to S in CYS | https://www.rcsb.org/ligand/CSX |
| HIC | add CH3 to HIS | https://www.rcsb.org/ligand/HIC |
| HYP | add OH to PRO | https://www.rcsb.org/ligand/HYP |
| YCM | add C-CONH2 to CYS | https://www.rcsb.org/ligand/YCM |
| YOF | add F to TYR | https://www.rcsb.org/ligand/YOF |
| M3L | add 3 methyl groups to LYS | https://www.rcsb.org/ligand/M3L |
| PFF | replace OH with F in TYR | https://www.rcsb.org/ligand/PFF |
| CGU | add COO to the second carbon in GLU | https://www.rcsb.org/ligand/CGU |
| FTR | add F to TRP | https://www.rcsb.org/ligand/FTR |
| LLP | added the complex group to LYS | https://www.rcsb.org/ligand/LLP |
| SNN | cannot form continuing peptide bonds | https://www.rcsb.org/ligand/SNN |
| CAF | add DIMETHYLARSINOYL to CYS | https://www.rcsb.org/ligand/CAF |
| CMH | add Hg-met to CYS | https://www.rcsb.org/ligand/CMH |
| MHO | add O to S in MET | https://www.rcsb.org/ligand/MHO |

### Multiple sequence alignments

There is no one-letter code for these noncanonical amino acids, which means that alignments can’t be made. The only ones that can be mapped are:

SEC’:’U’, ‘PYL’:’O’, ‘GLX’:’X’,. However, since GLX maps to X, it will look like “UNK” (unknown), which also maps to “X”. In addition, SEC and PYL are not even among the top 50 AAs. Therefore, we substitute all non-std AAs with X for the MSA search. AlphaFold2 (and later versions) generates three different MSAs. This process constitutes the main bottleneck for the predictions, as very large databases such as the Big Fantastic Database [36,37] are searched, which is very time-consuming [38]. To simplify this process, we instead search only uniclust30_2018_08 [39] with HHblits (from HH-suite [23] version 3.1.0):

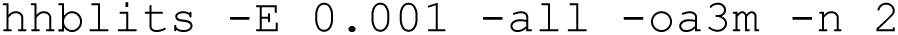

### Training and evaluation sets

To compare the method developed here (RareFold) with AlphaFold3 [20] (AF3), which can handle amino acid modifications, although these have to be specified on the atomic level, we analyse what structures in the PDB have less than 20% sequence identity to the AF3 training set (all data in the PDB up to 2021-09-30) and contain ncAAs. Before 2021-09-30, there were 62,530 examples, and after 12,352, belonging to 8125 and 2360 clusters, respectively, of which 906 are only found after the date cutoff. In these 906 clusters, there are 1654 unique structures, and 177 of these have modified amino acids. However, only 59 are non-MSE, leaving relatively little evaluation data for the other amino acid types.

In total, there are 7599 structures (10.1%) with modified amino acids, making the 177 not in the AF3 training set only 2.3% of the total set. To obtain a valid comparison, we choose to continue with this set and train on all examples before 2021-09-30, test on the 177, and validate on the remainder (Table 2). Supplementary Figure 8a displays the length distribution for the different partitions, and Supplementary Figure 8b the number of ncAA in each protein. For the test set, most proteins are of shorter lengths, while the training and validation sets correspond well to each other.

**Table 2.** Training, validation and test partitions.

| Partition | Examples | Modified examples | 20% clusters |
| --- | --- | --- | --- |
| Train | 62530 | 6679 (10 SNN) | 8125 |
| Valid | 12175 | 743 | 2265 |
| Test | 177 | 177 | 140 |
| Total | 74882 | 7599 | 10530 |

### Training

To bias the learning to noncanonical amino acids, we ensure that we always include examples with noncanonical amino acids in half of the batch, sampled according to the inverse frequency of the noncanonical amino acids. The other half of the batch is sampled according to the occurrence of the sequences in the 20% sequence identity clusters to enable the learning of diverse protein structures. We use a batch size of 24, take crops of 256 residues, and train across 8 A100 GPUs with 80GB of vRAM each. We maintain a constant learning rate of 10^-3^ throughout training after 1000 initial transition steps. Figure 7 shows the training curve for individual losses and metrics defined as in AlphaFold2 [1]. We use the same loss function as in AlphaFold2:

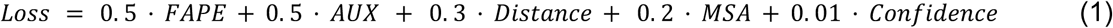

**Figure 7.**
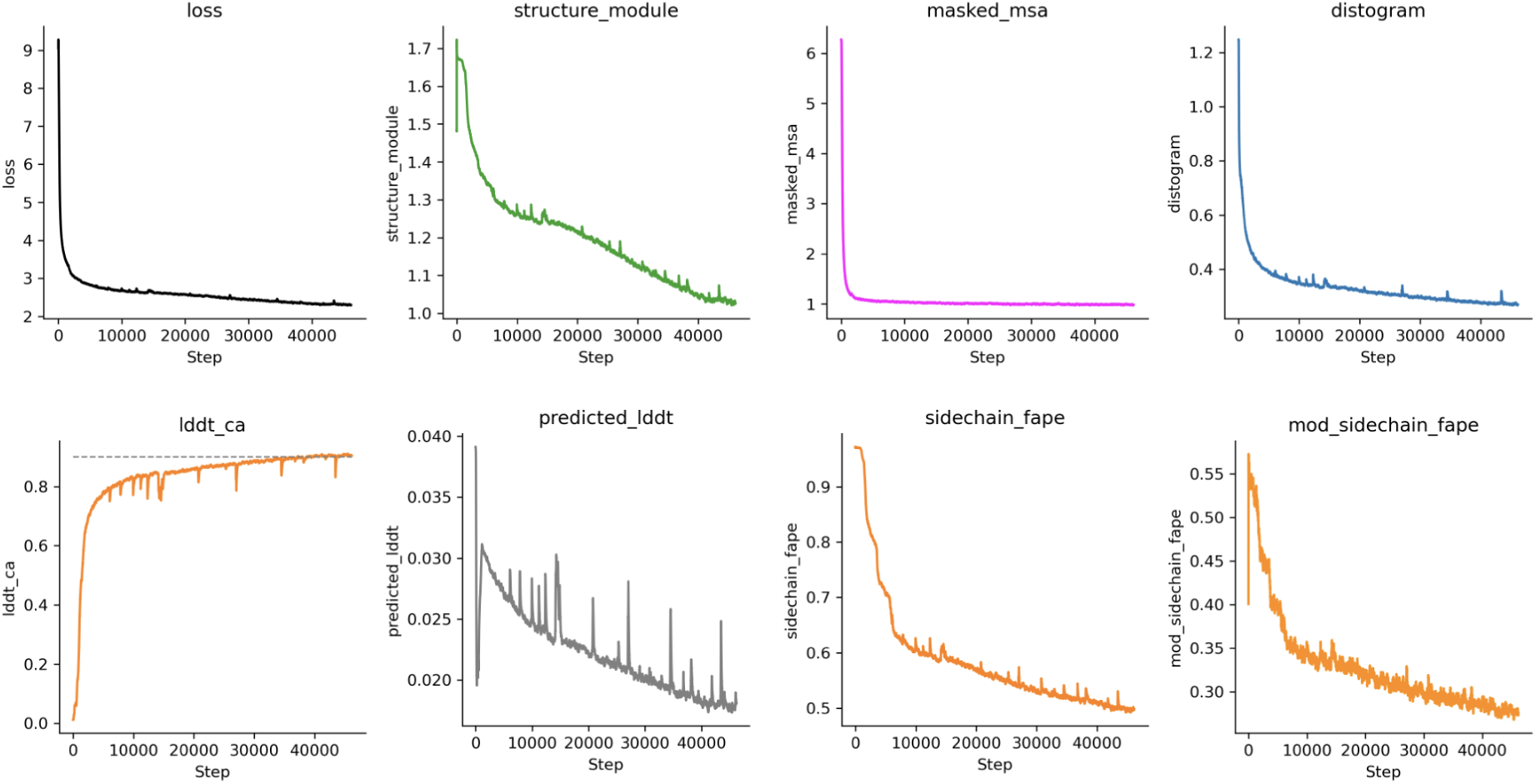
Training curves for the RareFold structure prediction model. Training losses and metrics over approximately 40000 steps are shown with a smoothing window of 500. The losses include: the combined loss function (equation 1), AUX (structure_module, a combination of FAPE and angular losses), distogram (pairwise distance loss), FAPE (frame aligned point error) for regular amino acid side chains and modified (mod_sidechain_fape), and predicted_lddt (difference between true and predicted lDDT scores), all defined as in AlphaFold2 [1]. The Cα lDDT is also monitored. From step 20000, the model undergoes fine-tuning with larger crops and additional losses applied to reduce residue bond violations, clashes, and extreme Cα–Cα distances (see Fine-tuning), resulting in improved structural quality without the need for relaxation.

Where *FAPE* is the frame aligned point error, *AUX* a combination of the *FAPE* and angular losses, *Distance* a pairwise distance loss, *MSA* a loss over predicting masked out MSA positions and *Confidence* the difference between true and predicted lDDT scores. These losses are defined exactly as in AlphaFold2, and we refer to the description there [1]. From step 20000, the model is fine-tuned further using larger crops and including additional losses (see Fine-tuning).

### Fine-tuning

To resolve clashes that may appear between residues, we apply the same fine-tuning losses with the same loss weights as used for AlphaFold2 [1]: between residue bond violations, clashes and extreme CA-CA distances. We do not fine-tune any intra-residue violations. We fine-tuned for 5000 steps, reaching a total of 25000 training steps using a constant learning rate of 10^-4^ ( Figure 8). We again train on 8 GPUs and now take crops of 384 residues, with 2 examples per GPU (16 examples per batch). The fine-tuning was performed mainly to obtain a model which does not require relaxation (as this is equivalent, see below). This model is useful for design purposes, where the extra relaxation step can become costly if performed at each iteration.

**Figure 8.**
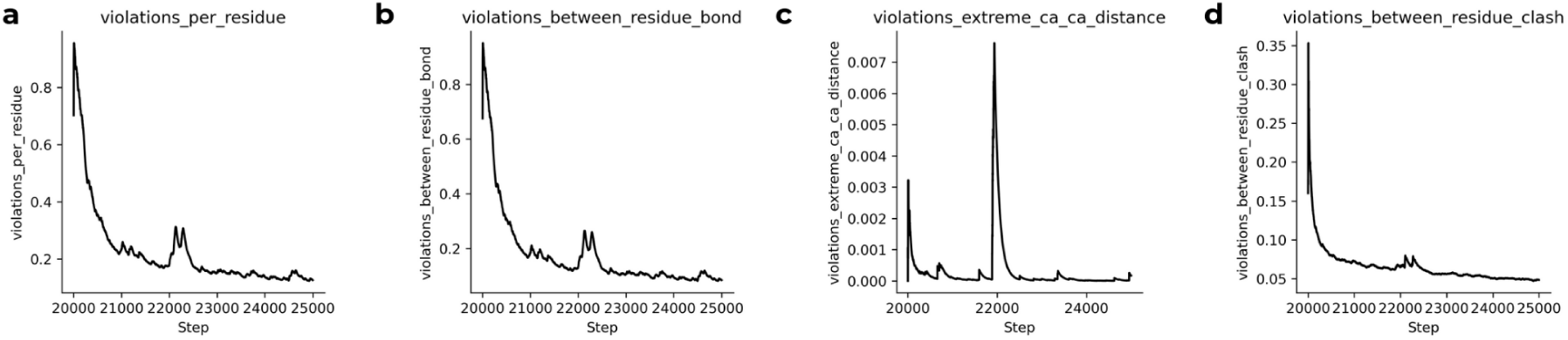
Fine-tuning improves structural violation metrics. Fine-tuning RareFold for 5,000 additional steps (total 25,000 training steps) using loss functions and weights adapted from AlphaFold2 [1] reduces residue **(a)** and bond violations **(b)**, extreme Cα–Cα distances **(c)** and clashes **(d)**. No intra-residue violations were fine-tuned. This process minimises structural clashes, producing models that do not require computationally expensive relaxation steps, enhancing efficiency for iterative design workflows.

To ensure RareFold scales to larger complexes and matches the accuracy of state-of-the-art atomistic models, we extended fine-tuning for an additional 25,000 steps (total 50,000 steps, Supplementary Figure 9) using larger crop sizes of 512 residues. Access to a cluster of 8 NVIDIA H200 GPUs (141 GB vRAM) enabled efficient training on these larger crops with a batch size of 16 (2 examples per GPU). This high-memory regime proved critical for modelling long-range interactions, ultimately raising the Global Cα-lDDT to parity with AlphaFold 3 (Figure 1).

### Relaxation

To resolve inter-residue clashes and improve local geometry, we apply molecular dynamics (MD) relaxation as a final step in structure prediction. We use OpenMM [40] with the CHARMM36 force field [41] and the Langevin middle integrator. Since ncAAs are not natively supported, we temporarily replace them with their closest canonical analogues during relaxation and substitute them back afterwards. While relaxation does not substantially alter the predicted structures, it slightly worsens most evaluation metrics (Figure 9). All comparisons between AF3 and RareFold were performed with unrelaxed structures, ensuring a strictly fair assessment.

**Figure 9.**
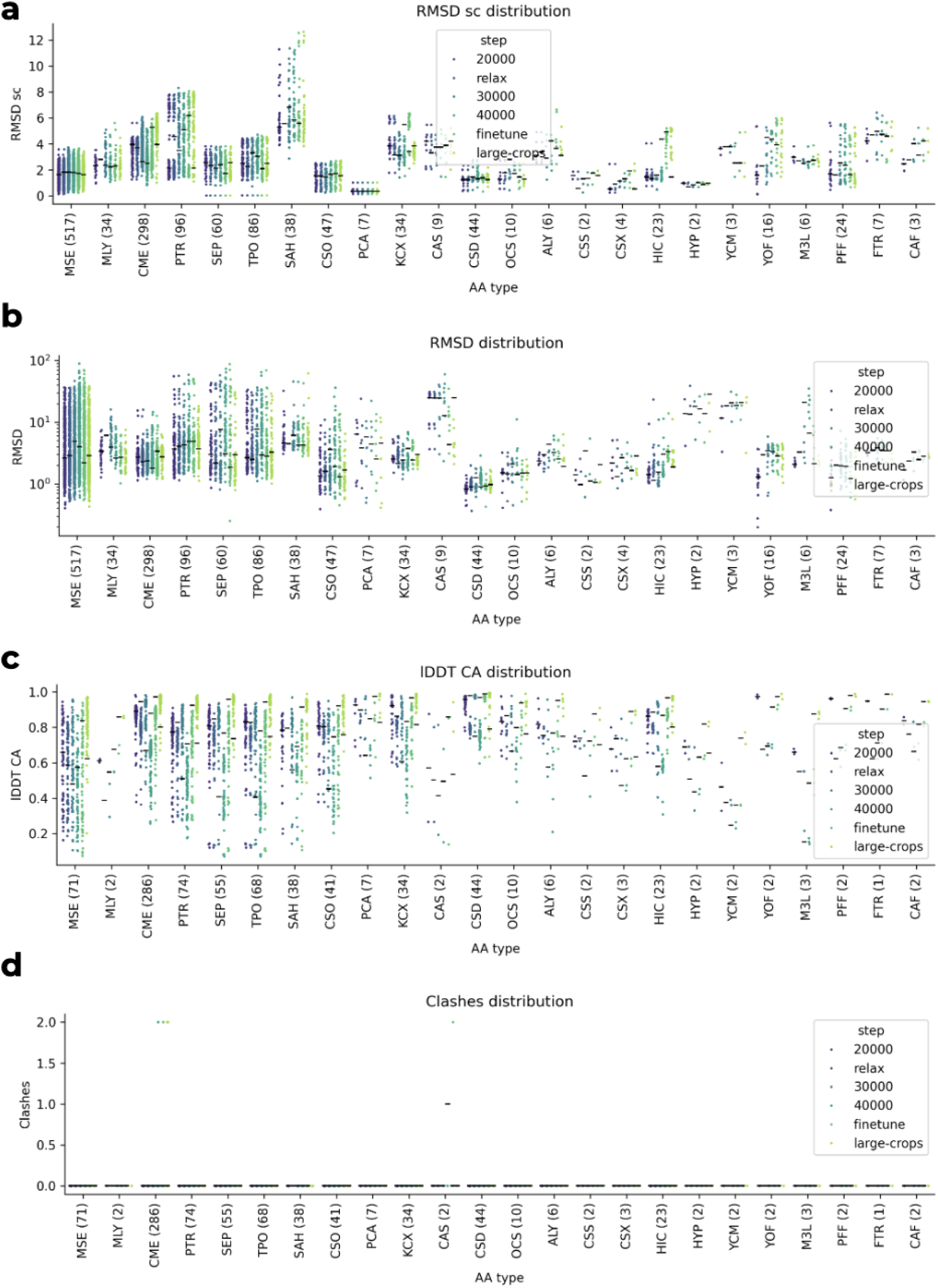
Validation metrics. The validation was performed at steps 20000, 30000 and 40000, with additional relaxation at step 20000, fine-tuning for 5000 steps from step 20000 (finetune), and from step 25000 for 25,000 steps with large crops (large-crops). Distributions are shown per ncAA for **a)** RMSD (Å) of side chains after side chain alignment, **b)** RMSD of side chains after global alignment, **c)** Overall Cα lDDT scores and **d)** intra-residue clashes. Internal clashes are only observed at 40,000 steps, finetune and large-crops for CME and for 40,000 steps and finetune at CAS.

### Validation

We validated the baseline model every 10,000 steps, starting at step 20,000 using the 743 examples with modified amino acids from Table 2. For each example, one structure was predicted using the previously generated HHblits MSAs and three recycles. Of these, 731 (98.3%) were successfully scored, with 12 excluded due to structural inconsistencies. Figure 9 shows that relaxation, performed at step 20,000, does not significantly impact accuracy. However, not all residues could be included in the relaxation due to missing representations in OpenMM, resulting in 679 (93%) structures being relaxed. The biggest effect comes from the large crop fine-tuning, which improves the overall Cα lDDT for all amino acid types. We also analyse the effect of recycling in Supplementary Figure 1, which we find has no meaningful effect on the performance.

### Scoring

For the evaluation, we align the Cα atoms of the predicted structures and calculate the RMSD (Å) of modified atoms for each modified amino acid (mod AA), considering all atoms in that AA (RMSD). The overall Cα lDDT score is computed for global structure accuracy [22] (lDDT CA). For each modified amino acid, the backbone atoms (N, Cα, C) are also aligned, and the side chain RMSD is calculated for the remaining atoms(RMSD sc). The test set consists of 177 structures, of which 174 could be scored (three contained structural inconsistencies), for RareFold and 171 for AF3 (due to three being OOM using 80 GB of vRAM), and the validation set of 731 structures for RareFold.

### Recycling

We analysed the effect of varying the number of recycles on the baseline model (20,000 training steps) using 3, 5, 10 and 20 recycling iterations. Interestingly, the number of recycles does not significantly improve the structure prediction of the ncAAs (Supplementary Figure 1). No internal AA clashes were observed at any of the investigated number of recycles, except for in one case for TPO at 20 recycles.

### AlphaFold3

AlphaFold3 (AF3) [20] can be used to introduce modifications to amino acids and thereby generate ncAAs by the addition of covalent modifications. In the AF3 publication, only 40 examples were evaluated, and these were from proteins that had less than 40% sequence similarity with their training set. The predictions were evaluated based on whether the aligned RMSD of the ncAAs were <2 Å. Although only around half of these examples were predicted with an RMSD < 2 Å, the evaluation is not rigorous due to the small number of examples, high sequence overlap and disregard of the type of ncAA.

We evaluate AF3 on the test set outlined above, which contains 14 different ncAA types (Figure 1). We use HHblits MSAs and no templates (the same conditions as for RareFold). We take the first prediction using the default settings of 5 predicted models; modelSeeds”: [1], “dialect”: “alphafold3”, “version”: 1. We introduce modifications as explained in the AF3 GitHub: https://github.com/google-deepmind/alphafold3/blob/main/docs/input.md. In total, 174/177 proteins could be predicted, and 171 scored. Two failed due to being out of memory using NVIDIA A100 GPUs with 80 GB vRAM. The remainder failed for unknown reasons, but will not significantly impact the results.

### Artefacts from diffusion for MSE in AF3

One limitation of AF3 is the presence of unnatural atomic representations introduced by its diffusion module, leading to overlapping atoms, particularly in selenomethionine (MSE), the most common ncAA in the PDB. While AF3 appears to perform well on MSE based on RMSD metrics, a closer examination reveals systematic structural artefacts. MSE residues frequently exhibit steric clashes (atoms closer than 1 Å apart) due to improper placement of the selenium atom, leading to artificially low RMSD values that do not reflect realistic atomic configurations (Supplementary Figure 2). While post-hoc molecular dynamics could theoretically resolve these clashes, such expensive refinement steps are impractical for high-throughput design frameworks. More fundamentally, the generation of such sterically impossible states indicates that the model has not learned the intrinsic physical constraints of ncAA side-chains, a failure that is particularly detrimental when designing novel interfaces where precise packing is required. The reliance on RMSD as a sole metric for evaluating the performance of ncAAs is problematic. In the AF3 paper [20], a threshold of 2 Å is used to assess prediction accuracy; however, this does not capture the overall structural quality of ncAAs as the median scRMSD is close to 1 Å for MSE (Figure 1d).

Similar issues were observed with secondary structure predictions, where AF3 tends to overproduce ordered regions and underrepresent disordered loops due to biases in its diffusion-based training process [20]. To mitigate such problems, AF3 incorporates fine-tuning on AF2 predictions. However, this approach is unlikely to be generalizable to ncAAs due to the limited availability of training data. As the diffusion-based model primarily minimises RMSD, it is possible to find suboptimal solutions. This issue is inherent to diffusion-based approaches, as the stochastic nature of denoising makes it difficult to direct the model toward uncommon data regimes such as ncAAs.

### Binder design by inversion of RareFold

Building on our previous work with EvoBind [9], we inverted RareFold to create EvoBindRare (Figure 3a). We constrain the modified amino acids to a set that is readily available for synthesis from our supplier GenScript (Table 3). We initialise a random sequence using the standard 20 AAs and these 12 ncAAs, and update and score each sequence using the loss function:

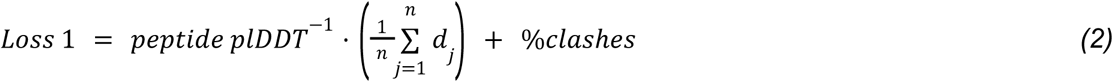

**Table 3.** Three-letter codes of amino acids available from GenScript (MSE, MLY, PTR, SEP, TPO, MLZ, ALY, HIC, HYP, M3L, PFF, MHO) and their corresponding names at GenScript.

| Three letter code | GenScript name |
| --- | --- |
| MSE | Se-Met |
| MLY | Lys(Me2) |
| PTR | pTYR |
| SEP | pSER |
| TPO | pTHR |
| MLZ | NMe-Lys |
| ALY | Lys-Ac |
| HIC | His(1-Me) |
| HYP | HYP |
| M3L | Lys(Me3) |
| PFF | L-4-F-Phe |
| MHO | Met(O) |

Where the peptide plDDT is the average plDDT over the peptide and dj is the shortest distance between all atoms n in the peptide and any atom in the target receptor. The %clashes is the number of protein-peptide atoms closer than 1.5 Å, divided by the number of peptide atoms. We run the design process for 1000 iterations using 24 different initialisations, lengths 10-20 and both linear and cyclic residue offsets [7] for the peptides.

For all design runs, three recycles were used. Due to a rewriting of the prediction with RareFold, we design in batches, resulting in the possibility to design 24 different binders simultaneously on a single NVIDIA A100 GPU with 40 GB of vRAM. The 1000 iterations are completed within 24 hours.

### Binder Design Selection

From the 1000 iterations x 24 initialisations x 11 lengths (n=264’000), we select the sequences with average peptide plDDT above 85 for both the linear and cyclic case. From these, we select the sequence for each length with the lowest loss (equation 2) for synthesis, resulting in 7 peptides for the linear case and 6 for the cyclic (Tables 4 and 5, Supplementary Figures 10 and 11). All linear peptides could be synthesised at a purity above 90%, but only one of the cyclic designs (length 14).

**Table 4.**
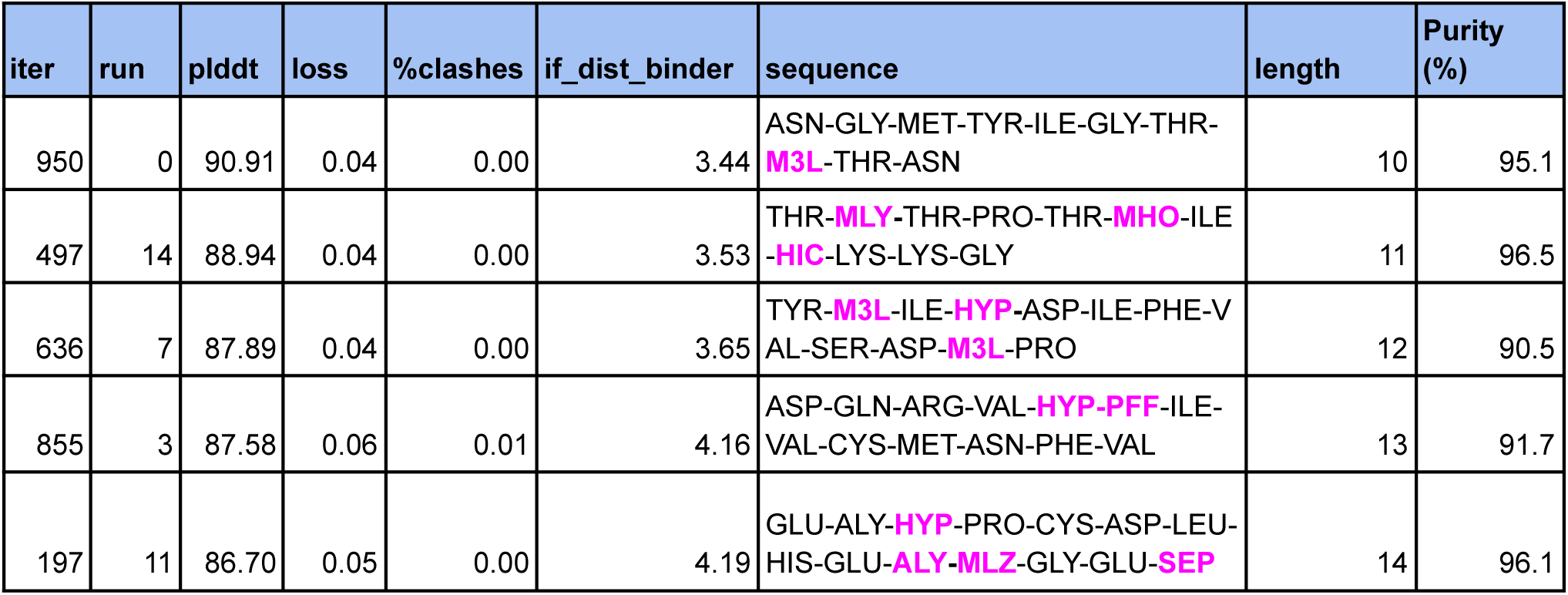
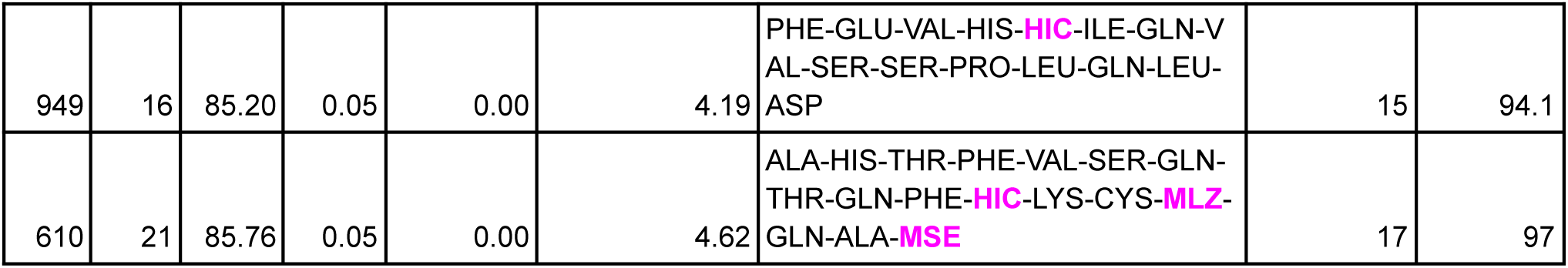
Linear selection (7 peptides). The table shows the iteration and run from which each peptide originated, predicted pLDDT (per-residue confidence), model loss, percentage of atomic clashes, interface distance to the binder, amino acid sequence, peptide length, and experimental purity as assessed by HPLC.

**Table 5.**
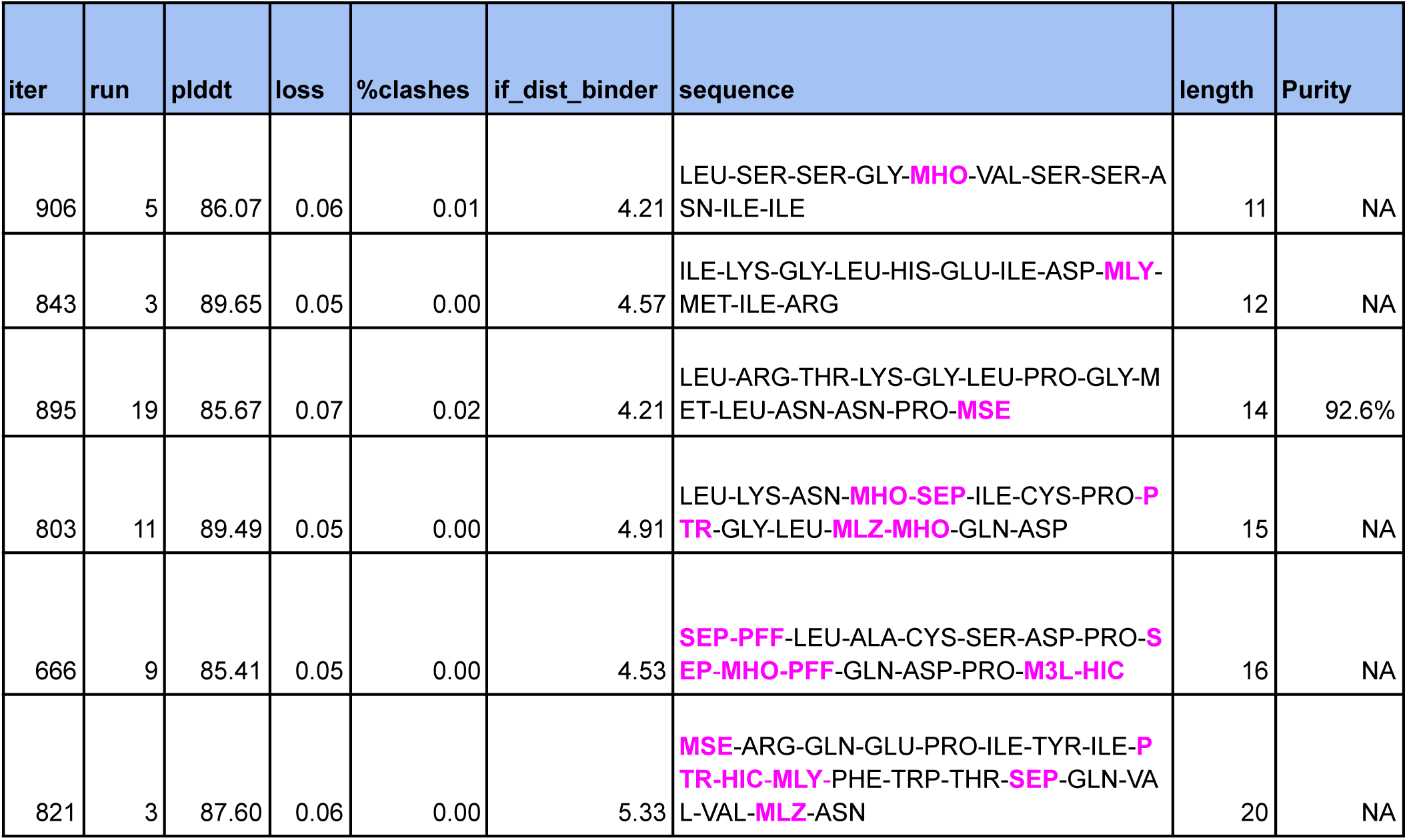
Cyclic selection (6 peptides). The table shows the iteration and run from which each peptide originated, predicted pLDDT (per-residue confidence), model loss, percentage of atomic clashes, interface distance to the binder, amino acid sequence, peptide length, and experimental purity as assessed by HPLC. Only length 14 could be synthesised at high purity (>90%).

### Efficient implementation reduces the computational footprint

To enable high-throughput design without prohibitive hardware costs, we implemented a consolidated batching strategy that maximises GPU utilisation. Standard implementations often require separate GPU processes for each sequence length or initialisation, for example, running 6 different lengths with 5 initialisations each would typically necessitate 30 separate GPU instances to execute in parallel. In contrast, our pipeline consolidates these varied inputs into a single batch on one GPU. For a typical design campaign (e.g., peptide lengths 10-15), this approach reduces hardware requirements by approximately 30-fold, allowing complex design trajectories to run on a single workstation (e.g., NVIDIA A100 40GB) within 24 hours. This stands in marked contrast to AlphaFold3, which lacks a native design framework and requires JIT recompilation for variable input sizes, creating significant bottlenecks during the iterative cycles required for design with ncAAs.

We further accelerated the design loop by parallelising non-inference tasks. The framework spawns multiple concurrent CPU instances to handle data processing, sequence mutation, and feature updates asynchronously. This strategy dramatically reduces “GPU idle time” (the latency between inference steps) from minutes to seconds. By overlapping CPU-intensive tasks with GPU inference, we achieved a massive reduction in wall-clock time per iteration (**Table 6**). The pipeline exposes parameters for sequence lengths and initialisations, allowing users to dynamically tune batch sizes to fully saturate available vRAM.

**Table 6.** Computational savings from efficient design implementation. Benchmarking was performed using the Ribonuclease (PDB ID: 1SSC) with binder length 10, the target previously validated in the design experiments here. Values represent the total resources and time required to complete a full design trajectory under standard versus optimised conditions.

| Design Step Component | Naive Implementation (30 Parallel Processes) | Optimised Batched Implementation (Single GPU) | Impact |
| --- | --- | --- | --- |
| Hardware Requirement | 30 GPUs | 1 GPU | 30 times Hardware Reduction |
| Prediction Time (GPU) | 91.3 s | 91.3 s | Equivalent throughput, vastly lower cost |
| Non-Prediction Overhead (CPU) | 600.6 s | 20.0 s | 97% Reduction in Overhead |
| Total Time per Step | ~692 s | 111.3 s | ~6.2 times Faster Wall-Clock Time |

### Peptide synthesis

Lyophilised powders of peptides designed by EvoBind2 were synthesised and purified by GenScript. The quality of successfully synthesised peptides (purity >90%) was verified by high-performance liquid chromatography (HPLC) and mass spectrometry (MS). The peptides were lyophilised with TFA salt and water, resulting in a net peptide content lower than the purity achieved before lyophilisation. The net peptide content was determined by GenScript, and serial dilutions for surface plasmon resonance measurements were calculated accordingly. Peptides were resuspended in HBS-EP+ buffer (10 mM HEPES, 150 mM NaCl, 3 mM EDTA, 0.005% Tween-20) to achieve a stock concentration of 1 mM.

Of the six cyclic peptide designs selected for experimental validation, only one (C14) was successfully synthesised and purified to the requisite quality (>90%). The remaining five designs failed during solid-phase synthesis or the subsequent cyclisation step. We attribute this attrition primarily to the inherent physicochemical properties of the specific full-length sequences, such as high hydrophobicity or aggregation propensities during the cyclisation reaction, rather than to a systematic incompatibility with the ncAAs themselves. This suggests that while the inclusion of ncAAs expands the design space, it also introduces additional complexity in the chemical synthesis of cyclic backbones that may require sequence-specific optimisation.

### Protein production

The expression of psfRNAseA-c001 was described previously [9]. Here is a brief description of the protocol: Transform the psfRNaseA-c001 construct (MBP-8xHis-TEV-RNaseA in pMAL-p2E plasmid) into *E. coli* BL21 (DE3) T1R pRARE2 cells. Inoculate 15 ml/L of Terrific Broth (TB) medium supplemented with 8 g/L glycerol, 50 µg/ml ampicillin, 34 µg/ml chloramphenicol, and 0.4% glucose. Grow overnight cultures at 37°C, 175 RPM. On day 2, cultivate 9 L of culture in the LEX system at 37°C until OD600 reaches 2, then reduce the temperature to 18°C. Induce protein expression at OD600 ∼3 with 0.5 mM IPTG and continue overnight at 18°C. Harvest cells by centrifugation (10 min, 4500 × g), resuspend in IMAC lysis buffer (1.5 ml/g cell pellet) with Complete EDTA-free protease inhibitor and benzonase nuclease, and freeze at −80°C. In purification, Thaw cell pellets, disrupt by pulsed sonication (4 s on/off, 4 min, 80% amplitude), and centrifuge (20 min, 49000 × g). Filter the supernatant through a 0.45 µm filter and load onto a 5 ml HisTrap HP column, followed by a 2 ml HisTrap HP column using ÄKTA Xpress. Wash with IMAC wash 1 (10 mM imidazole) and wash 2 (50 mM imidazole) buffers, then elute with IMAC elution buffer (500 mM imidazole). Perform size exclusion chromatography (SEC) on a HiLoad 16/60 Superdex 200 column using gel filtration buffer (20 mM HEPES, 300 mM NaCl, 10% glycerol, pH 7.5). Pool fractions containing MBP-RNaseA fusion protein (∼160 mg yield). Add TEV protease to pooled fractions at a 1:15 molar ratio and incubate overnight at 4°C. Verify cleavage by

SDS-PAGE (**Supplementary Figure 12a**). Add 20 mM imidazole to the reaction mixture and pass through two 2 ml HisTrap columns to remove the MBP-His tag. Further purify by passing over 1 ml of Amylose resin. Collect the flow-through, concentrate using a Vivaspin 5 kDa MWCO filter, and exchange buffer to storage buffer (20 mM HEPES, 300 mM NaCl, 10% glycerol, pH 7.5) using a PD-10 column (**Supplementary Figure 12b**). Measure final concentration (8.2 mg/ml), aliquot (50 µl/tube), flash-freeze in liquid nitrogen, and store at −80 °C.

### Surface plasmon resonance

To measure the binding affinity of designed peptides to 1SSC, we employed surface plasmon resonance (SPR) using a Biacore 8K system. SPR detects real-time biomolecular interactions by monitoring refractive index changes near a sensor surface. Experiments were conducted at 25°C in HBS-EP+ running buffer (10 mM HEPES, 150 mM NaCl, 3 mM EDTA, 0.005% Tween-20).

1SSC was immobilized on a Series S Sensor Chip CM5 to over 10,000 RU via amine coupling with 1-ethyl-3-(3-dimethylaminopropyl)carbodiimide (EDC) and N-hydroxysuccinimide (NHS). Peptides were captured on the chip, yielding a maximum response (Rmax) of 25–50 RU. Single-cycle kinetics experiments used peptide concentrations of 2 nM, 20 nM, 200 nM, 2,000 nM, and 20 µM, diluted in the running buffer. Each concentration series was injected at 30 µL/min without regeneration, with 120 s association and 1,800 s dissociation phases.

Raw data were analysed using Biacore Insight Evaluation Software. Sensorgrams underwent reference subtraction (using an activated/deactivated reference flow cell) and blank subtraction (median of consecutive buffer injections) to correct for non-specific binding and bulk effects. Processed data were fitted to a 1:1 binding model [42] to derive global kinetic parameters: association rate constant (ka), dissociation rate constant (kd), and equilibrium dissociation constant (KD).

### Hydrogen-Deuterium Exchange mass spectrometry

A summary of all experimental parameters and results for the HDX-MS can be found in Supplementary Table 1, following standard reporting [43]. The info is also available in a separate table in the PRIDE upload (see Data availability section) with even more info, e.g. peptide-level data. All chemicals were from Sigma Aldrich, pH measurements were made using a SevenCompact pH-meter equipped with an InLab Micro electrode (Mettler-Toledo), and a 4-point calibration (pH 2,4,7,10) was made before all measurements. A DHR-PAL system (Trajan) was used for HDX sample preparation, with the labelling tray at 20°C, quench tray at 1°C, valve chamber and pre-chiller at 4°C, and digestion chamber at 8°C.

Peptide-bound samples were prepared by mixing protein (0.5 μg/μL) with 100 μM peptide, 20 mM HEPES, 150 mM NaCl, 1 mM EDTA, pH=7.5 in a 1:1 ratio (v/v). For the apo state, the peptide volume was replaced with HEPES buffer. HDX labelling was initiated by adding 2 μl sample to 45 μl dHEPES prepared in D20, pH_(read)_ 7.1. Labelling times were 0, 30, 300, and 3000 s with three technical replicates. Samples were quenched with 45 μl of 4 M urea, 0.2 M TCEP, 1% TFA, and 90 μl was injected. An UltiMate 3000 UPLC system (Thermo Scientific) was used for subsequent online sample handling.

Automated valve switching passed the injected quenched sample over a 2.1 × 20 mm Nepenthesin-2/Pepsin mixed digestion column (AffiPro, CZ) at 100 μL/min 0.1% HCOOH for 3 min, trapping the resulting peptides on a 1 × 5 mm trap (PepMap 100, C18, 5 μm), then desalted peptides at 100 μL/min for 1 min. Peptides were eluted and resolved by a gradient from 5% to 45% mobile phase B (95:5:0.1 CH_3_CN/H_2_O/HCOOH over 15 min on a 1 × 50 mm reversed-phase analytical column, Hypersil GOLD, particle size 1.9 µm. Eluting peptides were analysed on a Tribrid Eclipse Orbitrap mass spectrometer (Thermo Scientific) with HESI-2 electrospray ion source.

Full MS scans were acquired at a resolution of 120000 with a standard AGC target, maximum injection time on auto and covering an m/z range of 350 to 2,000 m/z. For peptide identification purposes, DDA tandem MS was performed on undeuterated protein samples. At the end of the analytical gradient, the solvent transitioned to 90% mobile phase B for 5 min, and halfway through that time, the analytical and trapping columns were put into back-flow washing mode by automated valve switching. After re-equilibration of the analytical column at 5% mobile phase B, the injection cycle ended at 22 min.

### Peptide immunogenicity analysis

The buffy coat was obtained from the blood service at Karolinska University Hospital, collected from a healthy adult donor, in accordance with institutional guidelines. Blood samples were collected from three individuals. Whole tonsils were collected from one individual undergoing surgery for obstructive sleep apnoea. Informed consent was obtained from all participants, and the study was performed under the approval of the Swedish Ethical Review Authority.

PBMCs were isolated from the buffy coat by density gradient centrifugation using Lymphoprep (STEMCELL Technologies), and tonsil tissues were dissected to generate single-cell suspensions [44]. PBMCs were adjusted to 1 × 10⁶ cells per mL in RPMI medium supplemented with 10% FBS, 1% penicillin–streptomycin, and 10 mM HEPES. Tonsil cells were resuspended at 1 × 10⁶ cells/mL in complete medium (RPMI with GlutaMAX, 10% FBS, 1× non-essential amino acids, 1× sodium pyruvate, 1× penicillin–streptomycin, 1× normocin (InvivoGen), and 1× insulin/selenium/transferrin cocktail (Gibco) supplemented with 0.5 µg/mL BAFF (BioLegend #559608). Cells (200 μL/well) were seeded in 96-well plates with either WT, C14, or L17 (25, 2.5, or 0.25 μM, respectively). Positive controls included cell stimulation with the TLR7/TLR8 agonist R848 (5 μg/mL, InvivoGen) or wells pre-coated with anti-CD3 and anti-CD28 antibodies (1 μg/mL each, Invitrogen) plus 5 ng/mL IL-2 for PBMCs. An unstimulated negative control was included. Cells were cultured in duplicate for 7 days at 37 °C in 5% CO₂ under humidified conditions.

Levels of proinflammatory cytokines (IL-6, TNF-α, IL-1β, IFN-γ, and IL-12) in culture supernatants collected on days 2, 5, and 7 were measured using an Invitrogen uncoated ELISA kit (Thermo Fisher Scientific, Vienna, Austria), according to the manufacturer’s instructions. The reaction was stopped, and absorbance was measured at 450 nm using a BioTek Synergy 2 microplate reader. Cytokine concentrations were calculated from standard curves generated using recombinant cytokines at known concentrations.

Anti-peptide antibody levels were evaluated by ELISA. 384-well plates (ThermoFisher Scientific, #464718) were coated overnight at 4°C with WT, C14, or L17 peptides (10 µg/mL). After washing with PBS containing 0.05% Tween-20, wells were blocked with 5% skim milk containing 0.1% Tween-20 in PBS. Culture supernatant collected on day 7, diluted 1:2 in 5% skim milk with 0.1% Tween-20 in PBS, or pooled serum from three individuals, diluted 1:20–1:640 in 5% skim milk with 0.1% Tween-20 in PBS, was added and incubated for 1.5 h at room temperature. After washing, horseradish peroxidase-conjugated goat anti-human IgG (Invitrogen, #A18805), IgM (Invitrogen, #A18835), or IgA (Jackson ImmunoResearch, #109-036-011), diluted 1:5,000 in 5% skim milk with 0.1% Tween-20 in PBS, was applied for 1 h at room temperature. Bound antibodies were detected with tetramethylbenzidine substrate (Sigma, #T0440), and the reaction was stopped with 0.5 M H₂SO₄ after 15-30 min. Absorbance was measured at 450 nm.

Cell phenotype was analysed by flow cytometry on day 7. Cells were filtered through a 70-μm cell strainer and washed with PBS. Cells were stained in three consecutive steps: first cells were incubated with live/dead Aqua (1:1000, ThermoScientific) for 20 min, then Fc-blocked (BD Biosciences, 1:100) for 10 min, and finally stained with a mix of the following anti-human antibodies, all from BD Biosciences unless otherwise stated: PE-Cy7 (CD19, 1:100), APC-Fire 810 (CD20, 1/100), APC-Cy 7 (CD3, 1/50), BV480 (CD4, 1/100), BUV496 (CD8 alpha chain, 1:100), BUV737 (CD27, 1:100), BUV421 (IgD, 1:200), BUV395 (IgM, 1:50), BUV510 (IgG, 1:200), PE (IgA, 1:2000), BUV805 (CD45, 1:50), BUV563 (CD16, 1:100), RB545 (CD14, 1:50), BV605 (TCRγδ, 1:50). All data were acquired on the SONY ID7000™ Spectral Cell Analyzer and analysed using FlowJo v10.9.0 (BD Life Sciences). Gating strategies for flow cytometry are provided in Supplementary Figure 13.

## Data Availability

All data presented here is available at https://zenodo.org/records/18420884 The raw HDX-MS data are available in ProteomeXchange via the PRIDE database with the project accession: PXD074453.

## Code Availability

The code for RareFold and EvoBindRare is available at https://github.com/patrickbryant1/RareFold.

## Acknowledgements

The protein purification was facilitated by the Protein Science Facility at Karolinska Institutet, Stockholm, and we would like to thank Dr Emilia Strandback, Dr Tom Reichenbach, Dr Henrik Spåhr, and Dr Tomas Nyman for assistance. We also thank Dr Likun Du and Dr Yating Wang, Division of Immunology, Department of Medical Biochemistry and Biophysics, at Karolinska Institutet, for their assistance with the flow cytometry experiments and data analysis. PyMOL and Blender were used for structural visualisation. We gratefully acknowledge Dr Simon Ekström and the Swedish National Infrastructure for Biological Mass Spectrometry (BioMS), the SciLifeLab Integrated Structural Biology Platform for providing facilities and experimental support. NIH Bioart was used to visualise the workflow in Figure 5a.

## Contributions

P.B. conceived and designed the study, developed RareFold and EvoBindRare, generated the binder designs, and wrote the initial manuscript draft. Q.L. performed the SPR affinity measurements. D.D. carried out structural visualisation in coordination with P.B., Q.L.. F.Z., H.M., and Q.P-H. performed the immune analyses. All authors contributed to manuscript writing and revisions, leading to the final version.

## Supplementary Information

### Supplementary Tables

**Supplementary Table 1.**
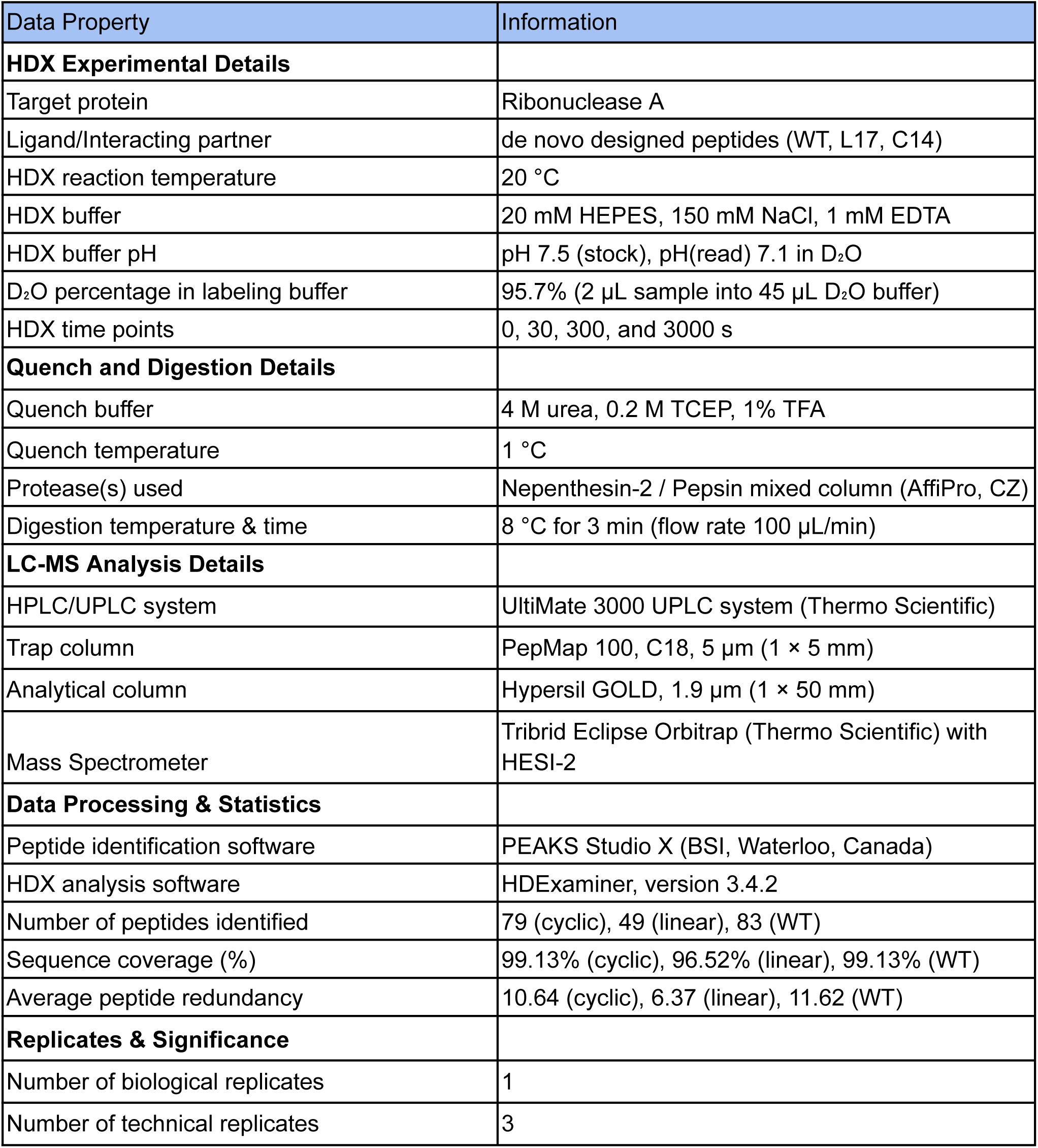
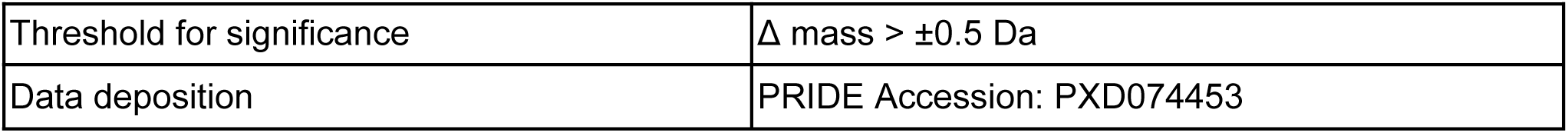
HDX-MS Summary.

### Supplementary Figures

**Supplementary Figure 1.**
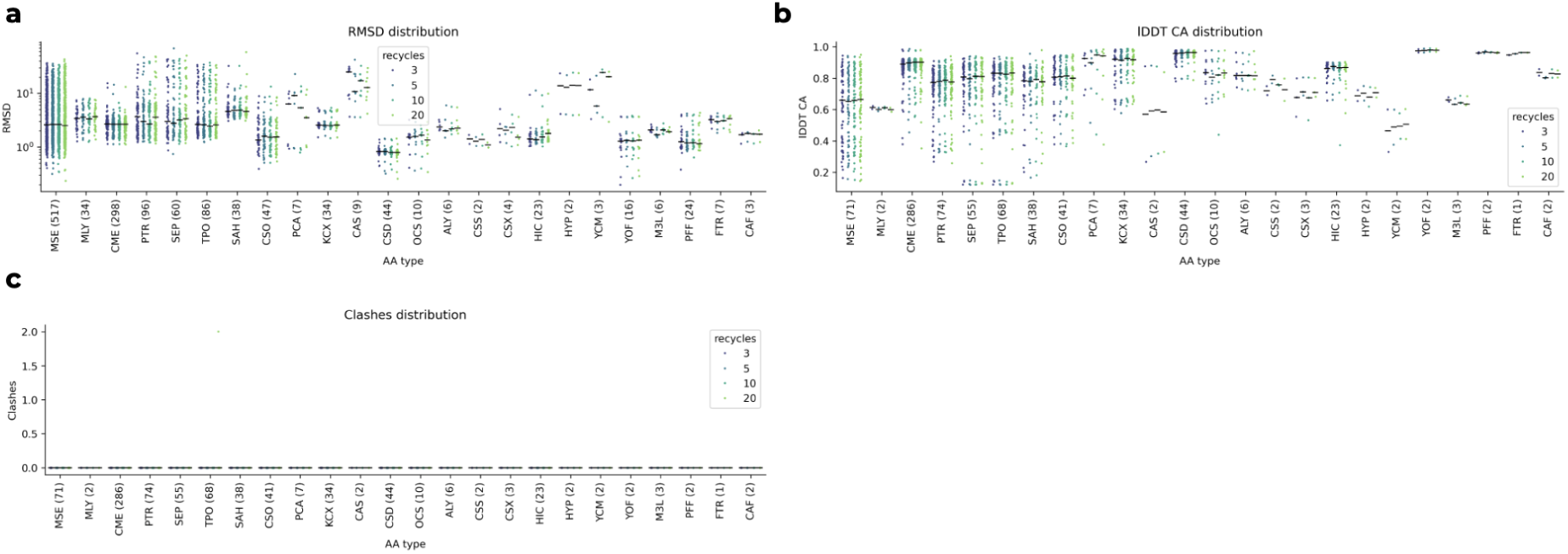
Number of recycles and accuracy metrics on the validation set for ncAAs. RMSD (Å), lDDT CA, and clash distributions are shown across different numbers of recycling steps using the baseline model trained for 20,000 steps. While the overall structural quality remains stable, additional recycling steps beyond a certain point do not lead to significant improvements in prediction accuracy. This suggests that RareFold converges early in the process, with ncAAs being predicted consistently even after fewer recycling steps.

**Supplementary Figure 2.**
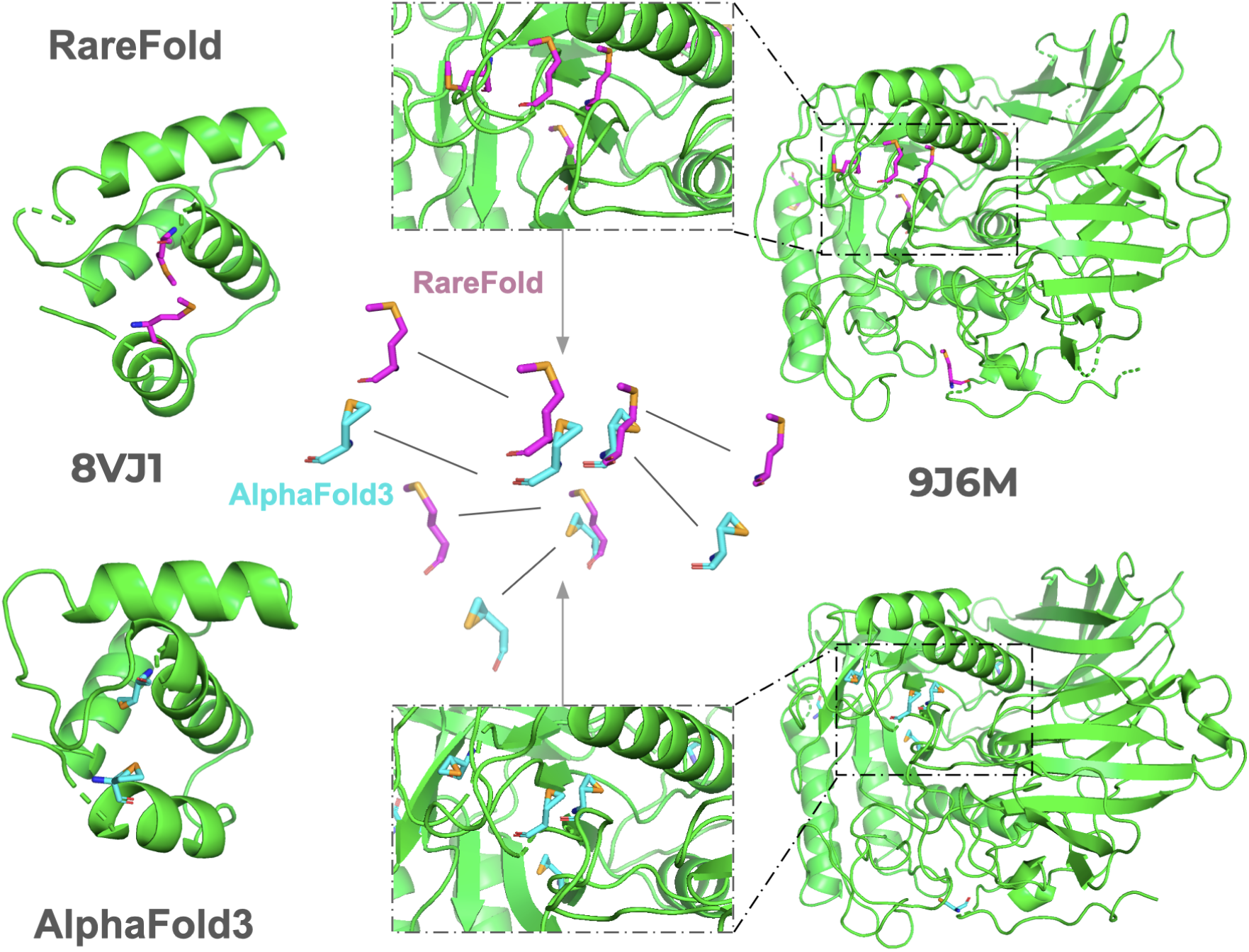
Limitations of AlphaFold3 in Predicting Modified Amino Acids. Comparison of MSE side chain predictions for two examples (PDB IDs 8VJ1 and 9J6M). The baseline RareFold model weights were used here, but the same results for MSE are found with the model fine-tuned on large crops (see the clash distributions in Supplementary Figure 3c). AF3 frequently produces unnatural atomic configurations for MSE (magenta sticks for AF3, cyan for RareFold) due to limitations in its diffusion module. These include steric clashes (atoms <1 Å apart) caused by misplacement of the selenium atom. Although AF3 reports good performance on MSE using RMSD, structural inspection reveals unfolded side chains and systematic artefacts not captured by this metric. The over-reliance on RMSD, including a 2 Å success threshold, fails to reflect true accuracy; median side chain RMSD for MSE is close to 1 Å (Figure 1d).

**Supplementary Figure 3.**
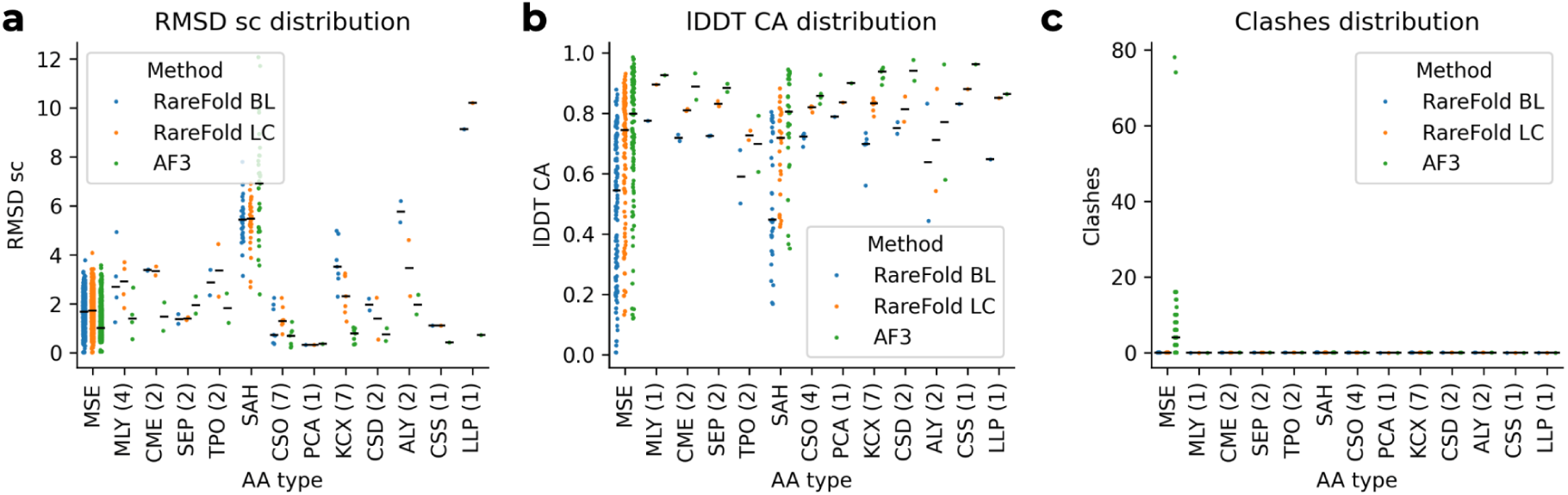
Test set performance of RareFold baseline versus fine-tuned models and AlphaFold3. Comparative analysis of structural accuracy on the held-out test set (n=174 for RareFold, n=171 for AF3). The RareFold Baseline represents the model performance before the final training stage. The Fine-tuned model (the final RareFold version) was further trained on 512-residue crops, allowing the model to learn longer-range contacts and improve global packing. This fine-tuning step yields consistent improvements in both global backbone geometry (Cα-lDDT) and side-chain accuracy (RMSD), bridging the gap towards the AlphaFold 3 benchmark while maintaining the inference efficiency of the token-based architecture.

**Supplementary Figure 4.**
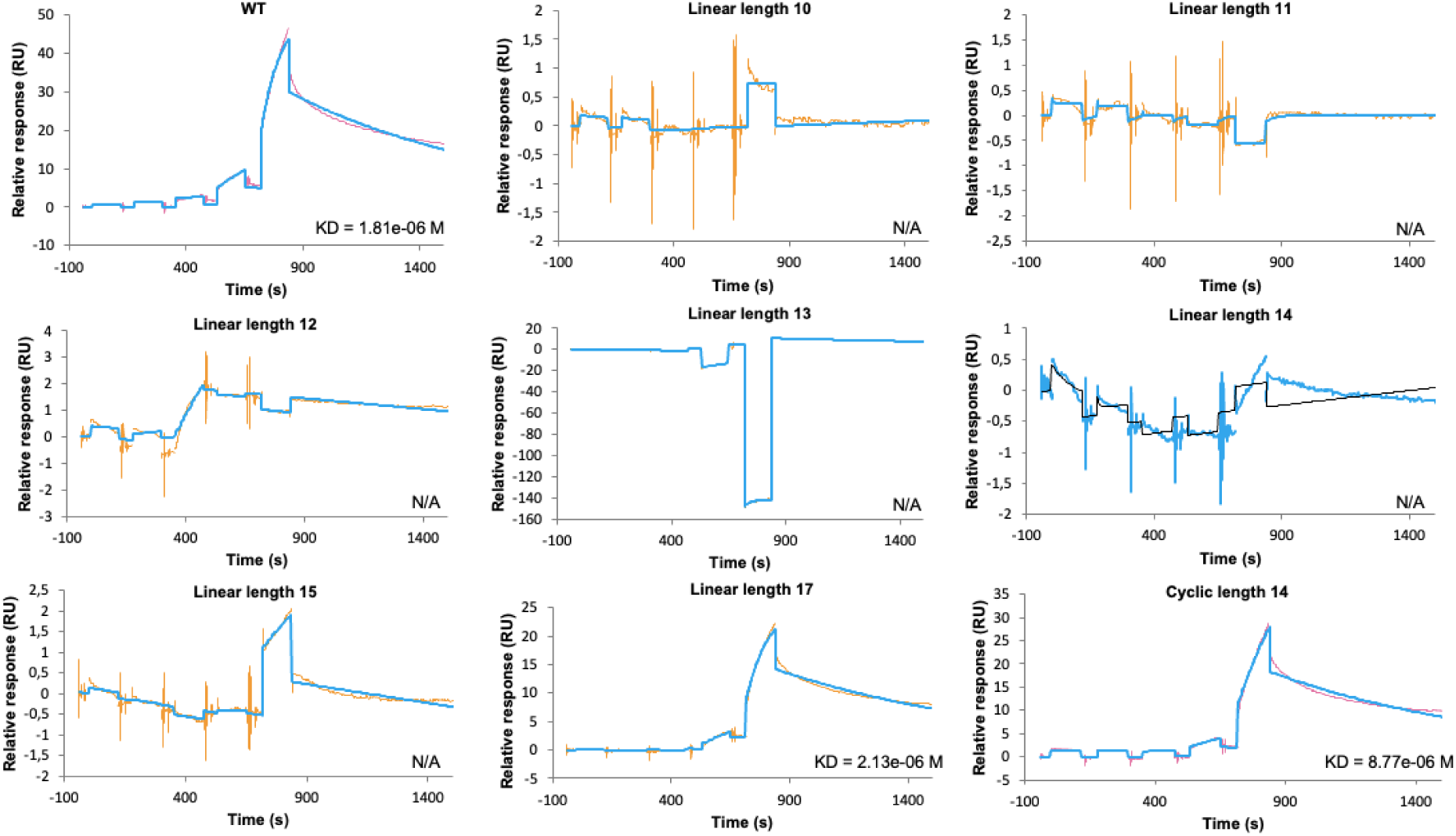
Sensorgrams for all evaluated peptides. Surface plasmon resonance (SPR) sensorgrams were generated using a Biacore 8K system. Each graph depicts real-time binding interactions between immobilised ligands and analytes in solution. Each subfigure represents a separate binding experiment, displaying the relative units (RU) on the y-axis versus time (seconds) on the x-axis. The colored lines represent the experimental data, while the blue lines show the fitted curves based on a 1:1 binding model (Methods).

**Supplementary Figure 5.**
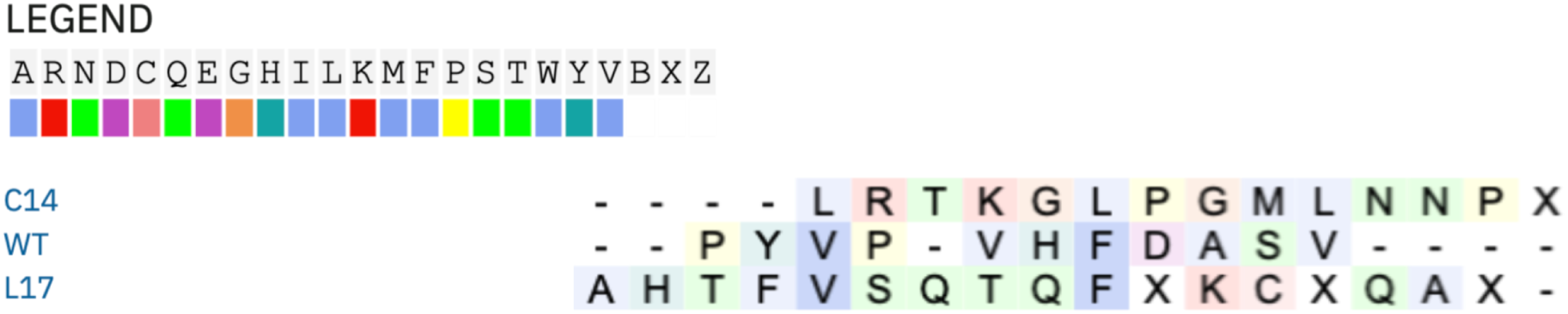
ClustalOmega (version 1.2.4) alignments of successful designs with the WT sequence. L17 has two matching residues with the WT (V and F), while C14 has none.

**Supplementary Figure 6.**
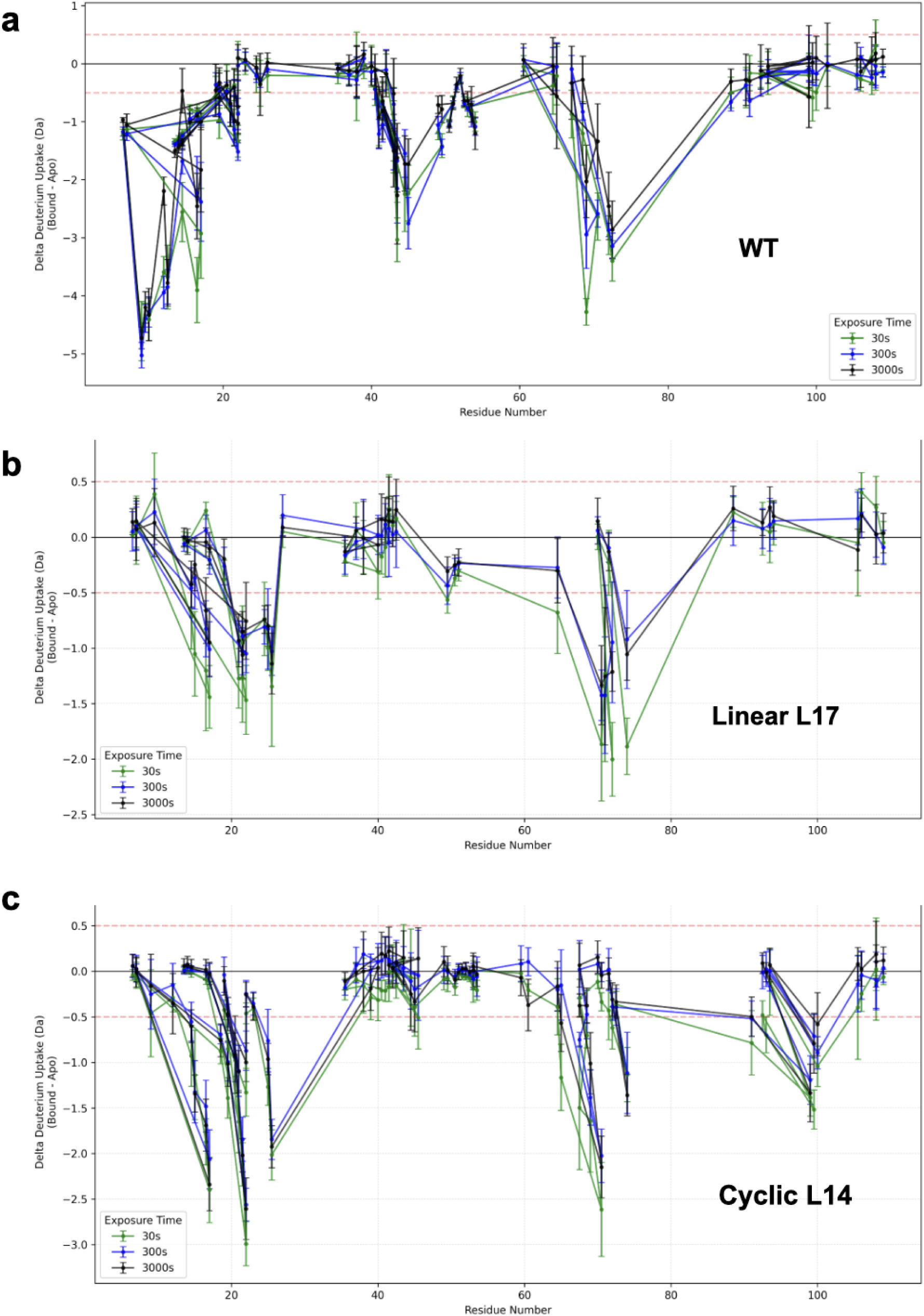
Differential deuterium uptake of the target protein upon binder engagement. The difference in deuterium uptake (Da) between the ligand-bound state and the unbound (apo) state is plotted against the residue number of the target protein. Data represents the sum of relative fractional uptake/uptake difference at 3, 300 and 3000 s for the target complexed with **(a)** the wild-type (WT) peptide, **(b)** the de novo linear design (L17), and **(c)** the de novo cyclic design (C14). Each bar represents a proteolytic peptide, mapped to its central residue. Negative values indicate regions of protection (reduced deuterium incorporation) corresponding to ligand binding or conformational stabilisation, while positive values indicate deprotection (increased solvent accessibility). The dotted line represents the significance threshold (0.5 Da); values outside this region are considered significant. Note that both L17 and C14 induce protection patterns in the same structural regions as the WT binder, confirming engagement at the predicted interface. The variation between the peptides signifies their varying binding dynamics, as expected from the predicted protein-peptide complex structures.

**Supplementary Figure 7.**
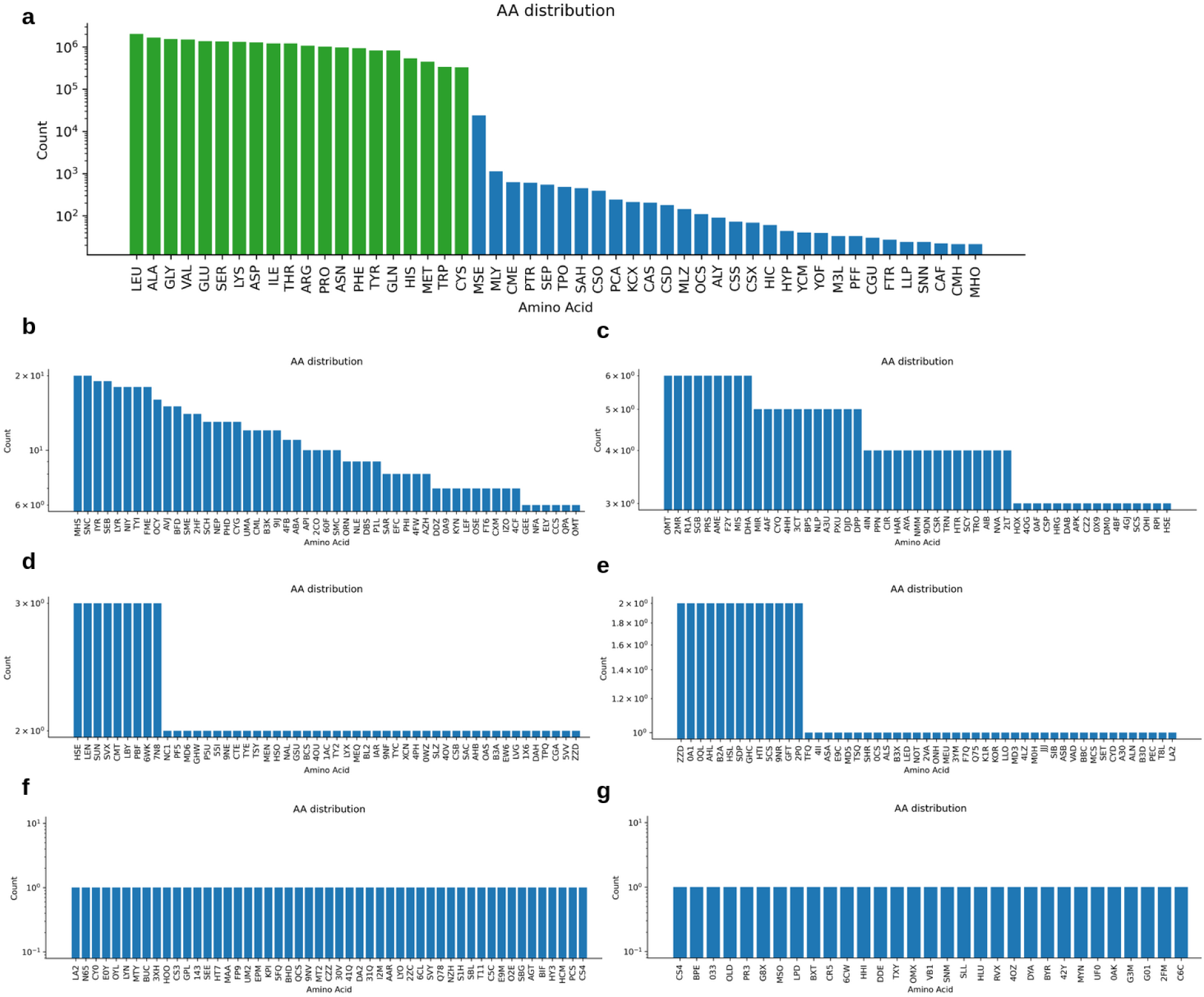
Amino acid distribution among single-chain structures in the PDB. **a)** Frequency distribution of the most prevalent amino acids, highlighting the 20 canonical residues (green) and the most frequent non-canonical amino acids (blue, e.g., MSE). **b–g)** Continued distribution of rare non-canonical amino acids ordered by decreasing frequency, illustrating the extensive “long tail” of chemical diversity present in the PDB. This dataset forms the basis for identifying the rare amino acids (ncAAs) used in RareFold training and evaluation.

**Supplementary Figure 8.**
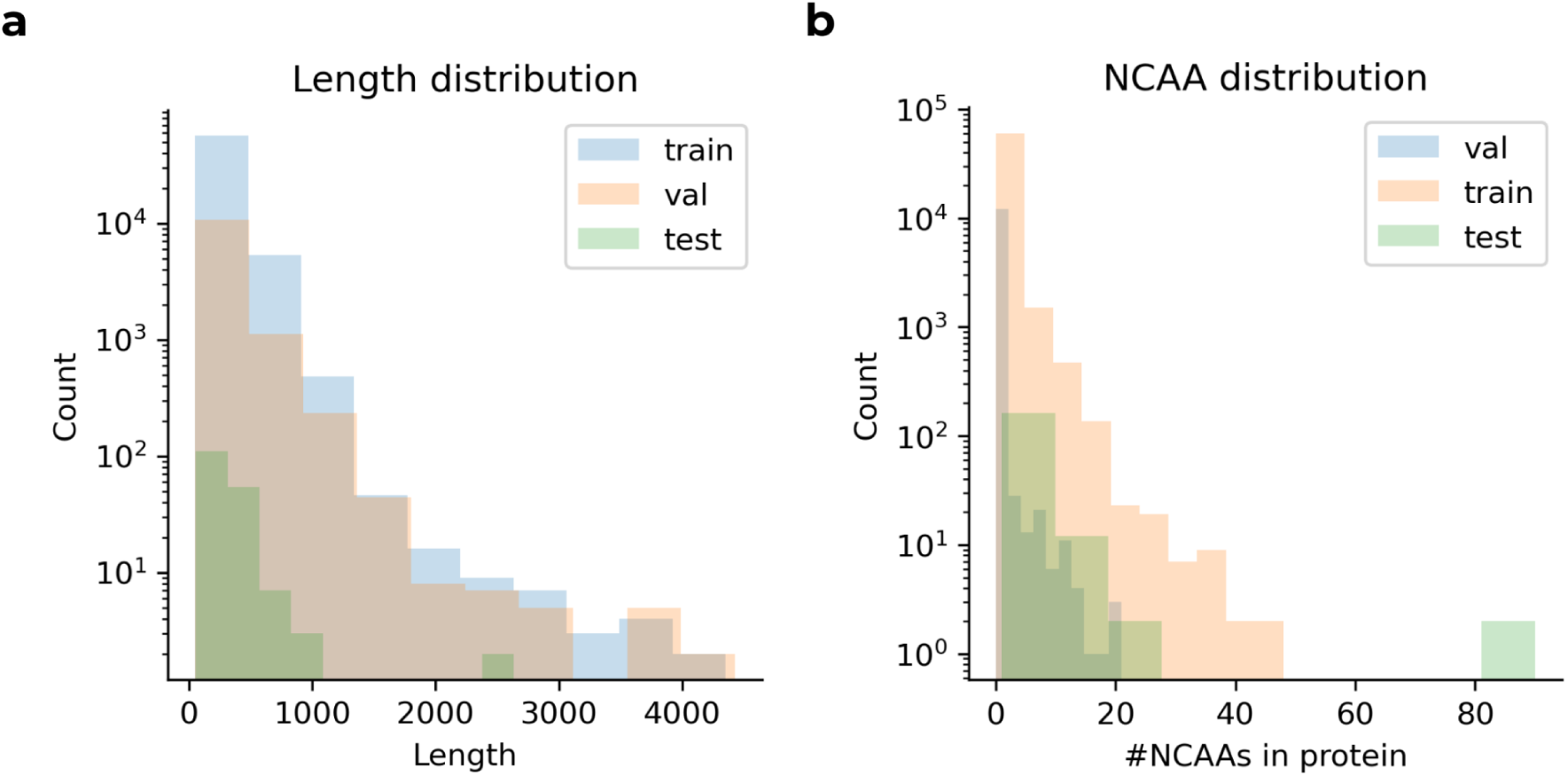
Length and modification distributions per partition. **a)** Chain length distribution per partition: The distribution of protein chain lengths in the training, validation, and test sets. **b)** Number of modified amino acids per protein and partition: The number of modified amino acids in each protein across the partitions. The training set contains more proteins with modifications than the validation and test sets, with an outlier in the test set featuring more than 80 modified amino acids.

**Supplementary Figure 9.**
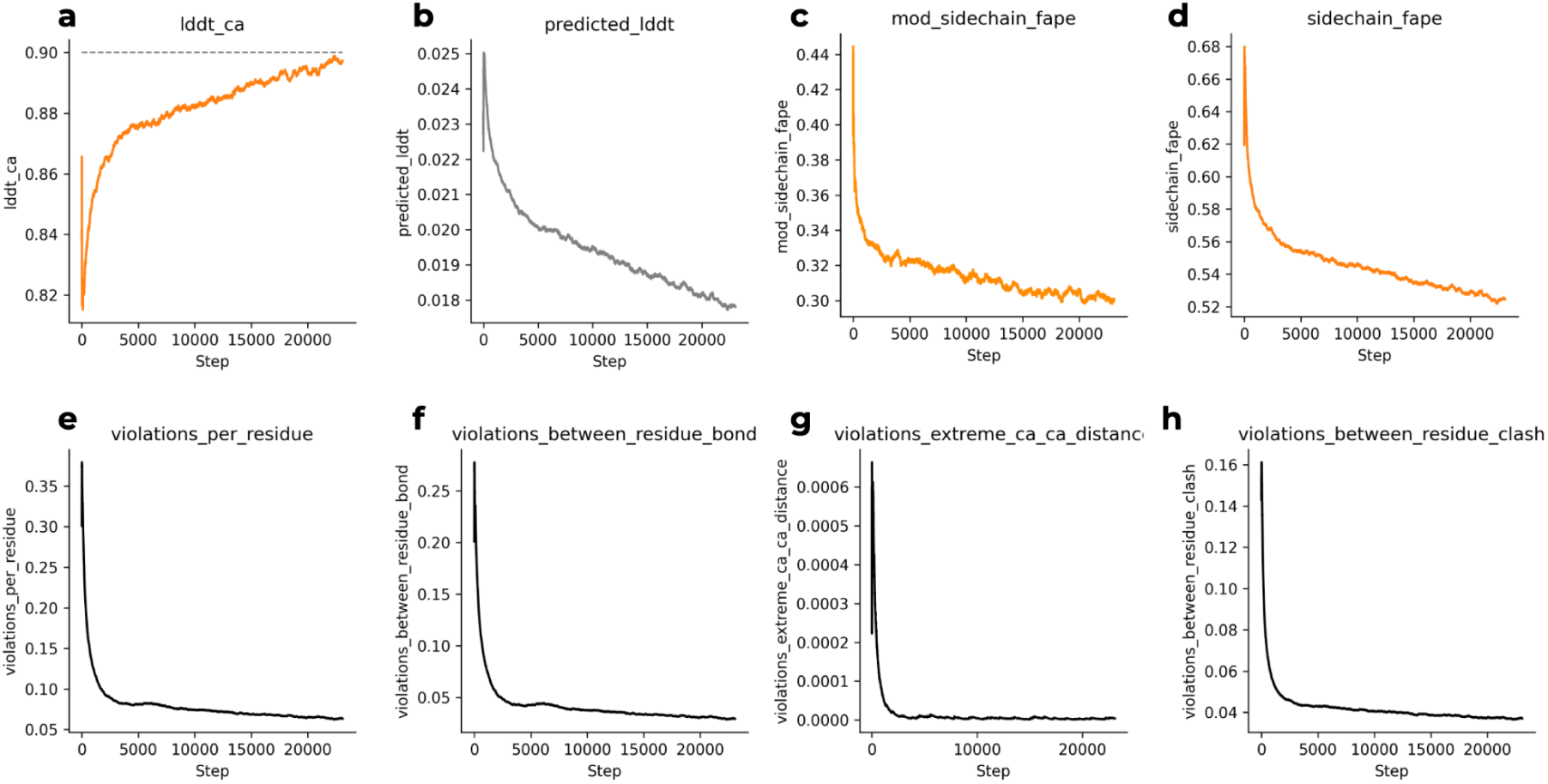
Fine-tuning on large crops (512 residues) for 25,000 steps. The model was fine-tuned to improve performance on larger complexes and modified residues. **a,** The Cα-lDDT score increases steadily, indicating improved global backbone accuracy. **b,** The predicted lDDT loss decreases, showing improved confidence estimation. **c, d,** Sidechain Frame Aligned Point Error (FAPE) decreases for both modified residues (**c**) and canonical residues (**d**), reflecting more accurate sidechain packing. **e–h,** Structural violation metrics show a rapid initial drop and sustained low levels for violations per residue (**e**), bond geometry (**f**), steric clashes (**h**), and extreme Cα-Cα distances (**g**), confirming the model learns to generate physically valid structures.

**Supplementary Figure 10.**
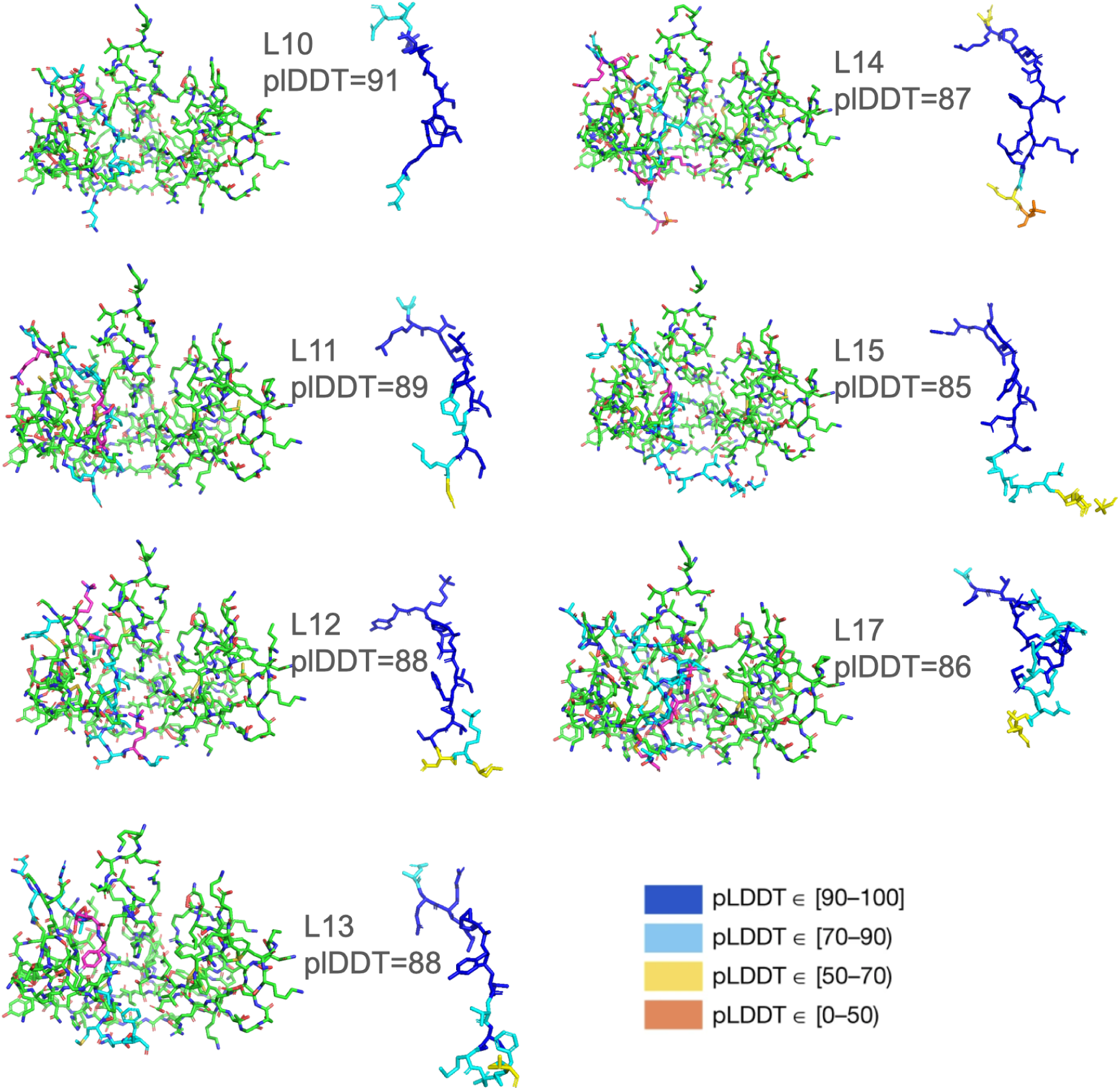
Linear selection of designed peptides (unrelaxed structures). Seven peptides were selected based on structural quality and interface metrics without applying structure relaxation.

**Supplementary Figure 11.**
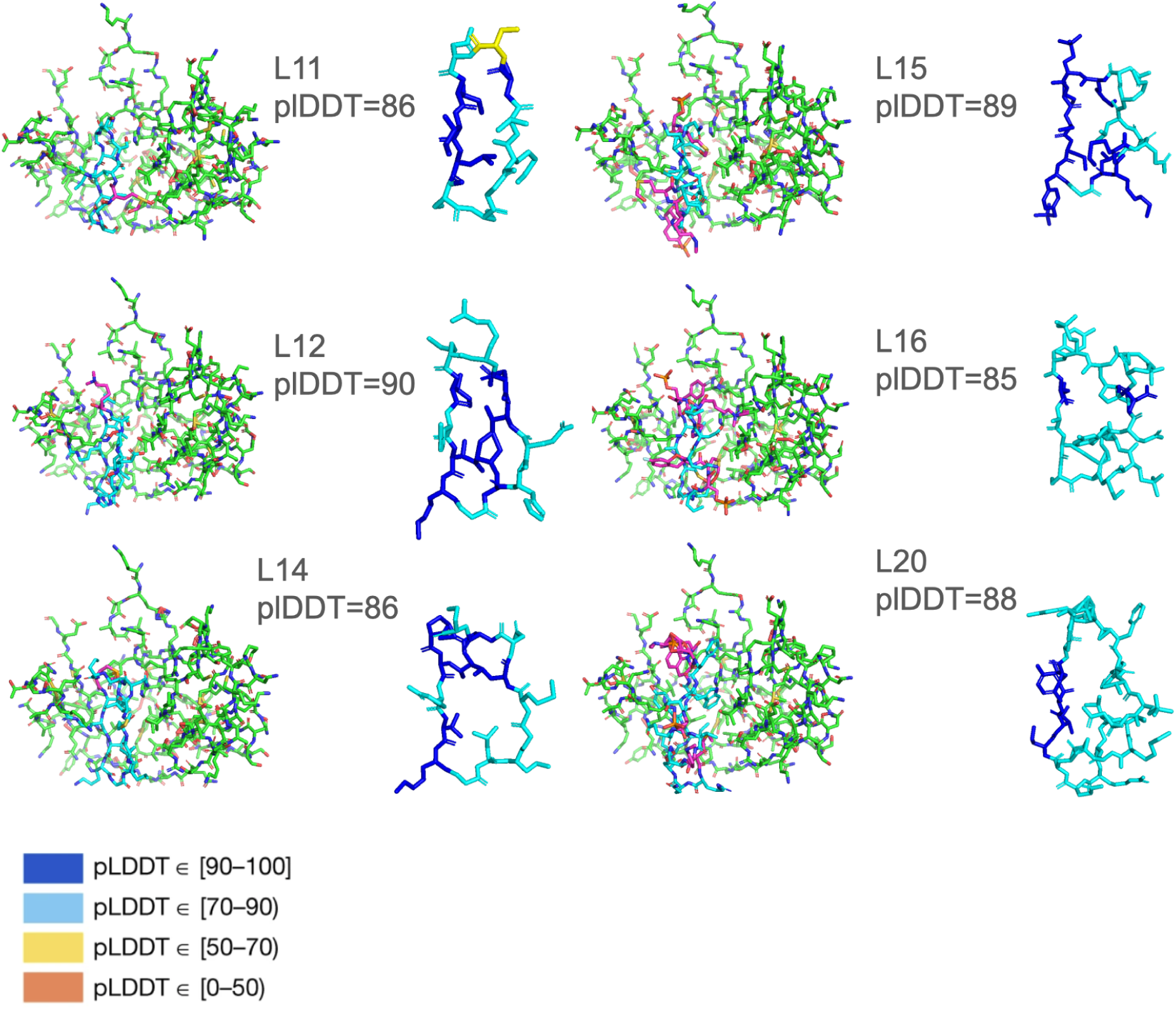
Cyclic selection of designed peptides (unrelaxed structures). Six peptides were selected based on structural quality and interface metrics without applying structure relaxation.

**Supplementary Figure 12.**
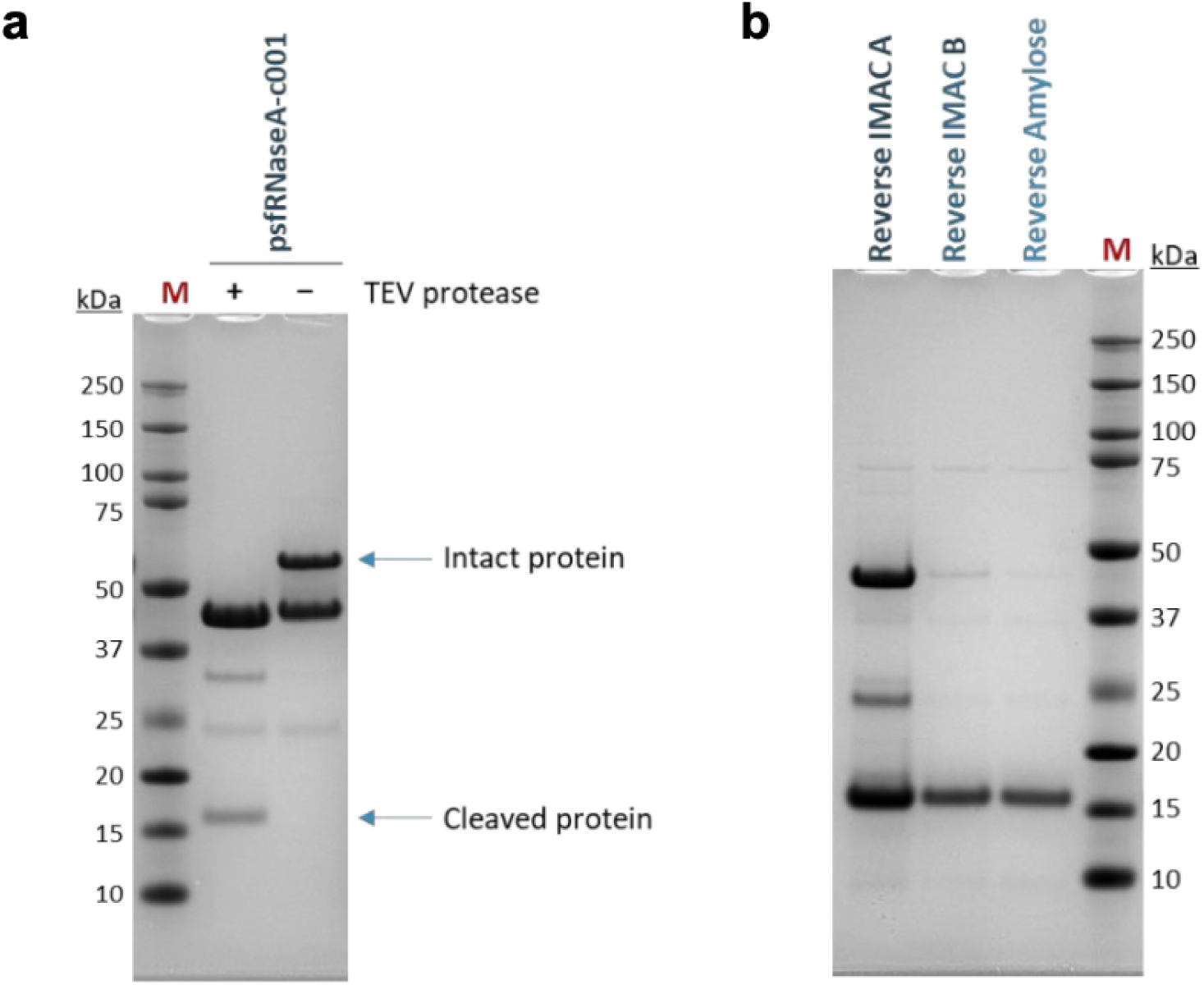
**a) SDS-PAGE analysis after tag removal with TEV protease**. TEV protease was added to the purified protein in a 1:15 molar ratio, and the reaction mixture was incubated in the cold room overnight. (+) indicates addition of TEV protease, (–) no addition of TEV protease. **b) SDS-PAGE analysis after reverse IMAC and reverse Amylose purifications.** The cleaved target protein was passed through two 2 ml HisTrap columns and one 1 ml Amylose resin to remove the MBP-His tag and TEV protease.

**Supplementary Figure 13.**
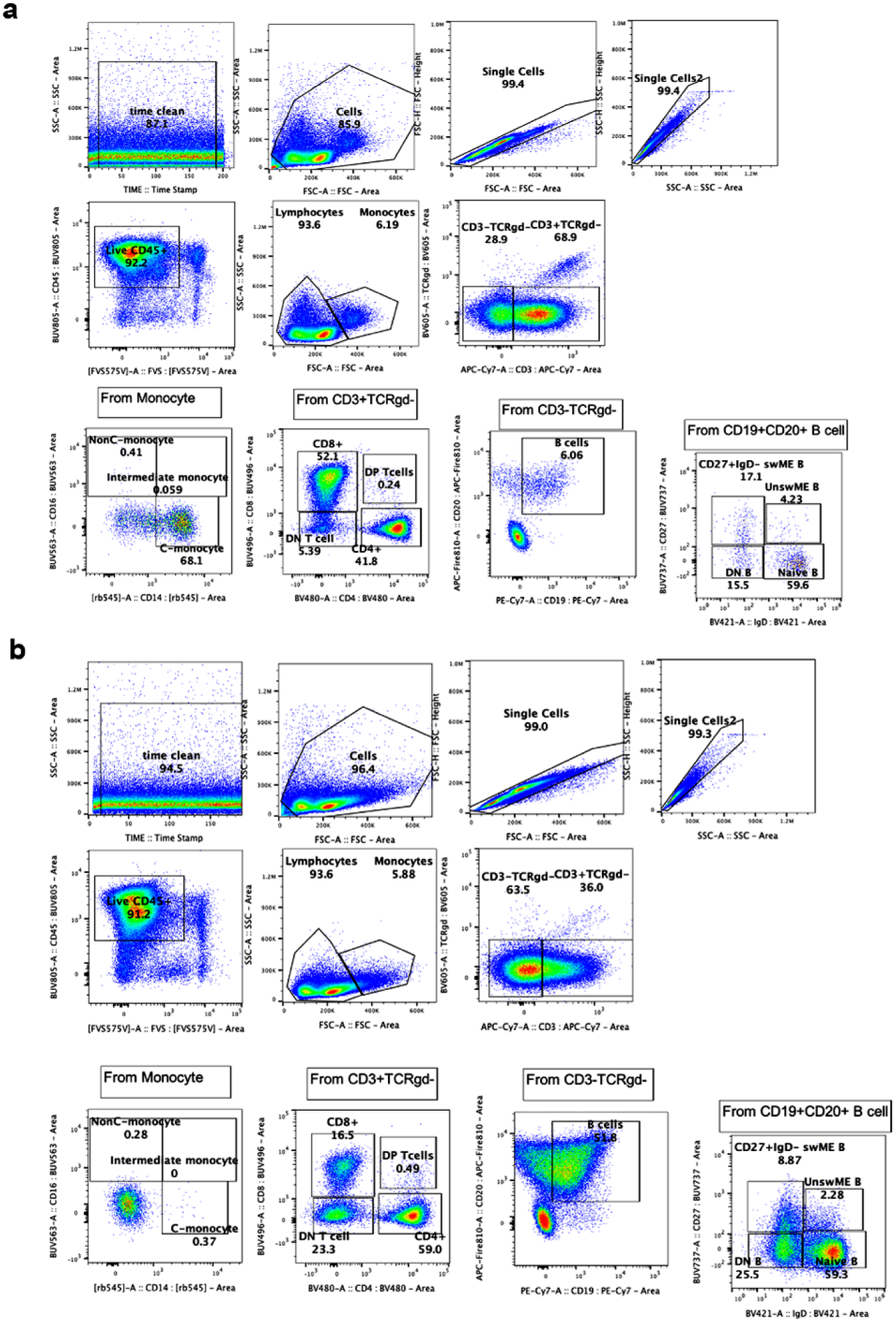
Representative flow cytometry gating strategies for lymphocyte and monocyte subsets. a) PBMCs, b) Tonsil cells. Flow cytometry plots show sequential gating of time clean, singlets, live CD45^+^ cells, separated into lymphocytes and monocytes based on their forward scatter (FSC) and side scatter (SSC). In the lymphocyte population, CD3^+^TCRγδ-cells were further gated into T cell subtypes: CD8^+^CD4^-^ (CD8 T cells), CD8^-^CD4^+^ (CD4 T cells), CD4^-^CD8^-^ (double negative (DN) T cells), D4^+^CD8^+^ (double positive (DP) T cells). CD19^+^CD20^+^ cells within the CD3^-^TCRγδ- gate were further subdivided into B cell subtypes: IgD^+^CD27^−^ (naïve B cells), IgD^−^CD27^+^ (switched memory (swME B) B cells), IgD^+^CD27^+^ (unswitched memory (UnswME B) B cells), IgD^−^CD27^−^ (double negative (DN) B cells). In the monocyte population, subtypes were further gated as follows: CD14^+^CD16^-^(classical (C-) monocytes), CD14^+^CD16^+^ (non-classical (NonC-) monocytes), CD14^-^CD16^-^(intermediate monocytes).

